# Task-Specific Computational Fluid Dynamics Evaluation of Multi-Outlet Extrusion Nozzles for Bioprinting

**DOI:** 10.64898/2026.01.09.698654

**Authors:** Cesar D. Vargas Urdaneta, Michael Taynnan Barros

## Abstract

Extrusion bioprinting is moving from proof-of-concept demonstrations toward repeatable, manufacturing-style workflows, but the nozzle remains a persistent bottleneck. Inside compact, opaque channels, bioinks experience strong geometric forcing that sets the local shear environment for cells, the pressure demand on the printer, and the uniformity of deposition. Multi-outlet nozzles are attractive because they promise higher throughput by splitting one feed into parallel filaments, yet in practice they often suffer from uneven outlet delivery and junction-driven shear hotspots. As a result, nozzle selection and scaling are still largely guided by trial-and-error rather than quantitative design evidence. In this work we provide a controlled, task-oriented comparison of two novel multi-outlet splitter archetypes— a compact radial 90° manifold and a branched Y-split—implemented with two and four outlets. Using three-dimensional computational fluid dynamics with representative rheology for three common hydrogel bioinks (GelMA, MeHA, and alginate) under typical pneumatic actuation, we resolve internal pressure, velocity, and wall shear stress fields and translate them into practical decision metrics for outlet balance and pressure-normalised throughput. The results show that the two-outlet 90° manifold is the most robust “default” geometry, consistently delivering the most uniform outlet splitting for shear-thinning inks while maintaining the most conservative shear footprint, making it the safest option for cell-laden and precision printing. The two-outlet Y-split achieves higher outlet speeds and is therefore better suited to fast deposition of acellular or support materials, but it concentrates elevated shear at the primary junction and is more sensitive to operating regime for weakly shear-dependent inks. Scaling from two to four outlets markedly increases the risk of maldistribution across all materials and pressures, and does not eliminate junction-anchored shear hotspots in the Y-split, indicating that passive geometric symmetry is insufficient at higher outlet counts. Overall, this study converts an informal design choice into evidence-based nozzle selection rules, enabling practitioners to match nozzle architecture to printing task (cell safety, precision, or throughput) and to anticipate when multi-outlet scaling will require active balancing or flow control. The proposed framework accelerates nozzle development, improves print reproducibility, and helps define safer operating windows for bioprinting with living cells.

## 1 Introduction

Three-dimensional (3D) bioprinting, the layer-by-layer deposition of biomaterials and living cells, has become an important technology for engineering tissues, organ models, and cell-laden constructs while providing precise control over the placement of cells and biomaterials at both millimetre and micro-scale levels (1; 2). A diverse toolbox of printing methods now enables the controlled placement of biomaterials and living cells with steadily improving control over the positioning of biomaterials and cells.(3). These methods include pneumatic and piston-driven extrusion, inkjet and drop-on-demand deposition, light-based printing (digital light processing or DLP, two-photon, and volumetric approaches), and embedded or freeform printing (extrusion into yield-stress support baths) (4; 5; 6; 7). The field is transitioning from proof-of-concepts to manufacturing-oriented workflows that balance print quality, viability, and repeatability against speed and cost (8; 9).

Extrusion bioprinting begins with digital design and material formulation, followed by the controlled deposition of bioinks during the printing phase, and concludes with post-printing maturation and assembly. This study focuses specifically on the printing stage, where nozzle geometry governs how applied pressure is transformed into internal velocity fields and wall shear stresses. These fluid-dynamic factors ultimately determine flow balance, print fidelity, and the mechanical environment experienced by embedded cells. Despite this progress, several challenges limit routine, high-quality bioprinting. Print fidelity can degrade through non-uniform flow, strand swelling, and die swell (extrudate expansion at the nozzle exit) (4). Cell viability can be reduced by excessive wall shear stress (WSS, the frictional force per unit area exerted by flow on channel walls) and extensional stresses (1; 2; 10). Practical operation must also respect pressure limits, footprint constraints, and requirements for sterilisable hardware (8). Many of these issues originate inside the nozzle, where complex, non-Newtonian bioinks experience geometry-dependent pressure drops and shear fields (11; 12; 13). However, computational fluid dynamics (CFD) studies that test different nozzle designs and operating conditions are still limited, so most optimisation relies on trial-and-error and leaves little basis for making informed design decisions (12; 14).

Nozzle geometry is a major design parameter that influences cell-compatible flow conditions and process performance through factors such as branching angle, curvature, taper, and outlet count (15; 16). Because in-nozzle phenomena (pressure, velocity, and shear-stress fields) are difficult to measure experimentally, CFD is a practical tool to evaluate potential nozzle designs before prototyping (17; 15; 18). With appropriate rheological models for shear-thinning bioinks and calibrated boundary conditions, simulations can map pressure, velocity, and WSS fields, quantify outlet flow balance (uniformity of discharge rates across multiple nozzles), and predict safe operating windows that are otherwise costly to explore experimentally (19; 20).

Designing extrusion nozzles requires balancing competing requirements that depend on the intended printing task. Geometric choices (branch angles, curvature, taper, outlet count, and path-length symmetry), operating conditions (pressure and flow control), bioink rheology, and printer integration constraints (sterilisability and footprint) interact in non-trivial ways to determine in-nozzle flow behaviour and, ultimately, printing performance (19). Within this design space, multi-outlet architectures are attractive because they can increase throughput by splitting a single inlet into multiple, parallel deposition streams. However, increasing outlet number can also amplify outlet-to-outlet variability and flow imbalance (17; 15). Likewise, geometric smoothing intended to reduce hydraulic losses can redistribute stresses and create elevated shear hotspots near junctions (20). These trade-offs make it unlikely that a single “best” nozzle geometry exists across applications. Instead, nozzle design should be *task-specific*, prioritising different objectives for cell-laden printing (minimising wall shear stress and maintaining uniform outlet flow) versus acellular, high-throughput deposition (maximising speed under acceptable shear and pressure limits). Multi-outlet nozzles therefore require a principled framework to determine when added outlets improve performance for a given task, and how the internal branching should be configured to control balance, shear exposure, and pressure drop. CFD provides this framework by resolving otherwise inaccessible in-nozzle fields (pressure, velocity, and wall shear) for realistic non-Newtonian inks and enabling controlled parameter sweeps over geometry, rheology, and operating pressure. Importantly, CFD quantifies task-relevant trade-offs (flow balance, shear hotspots, and pressure losses) with reproducible metrics before fabrication, allowing unsuitable designs to be eliminated early and safe operating windows for cell-laden extrusion to be bounded. This simulation-first approach focuses manufacturing and bench testing on a small set of evidence-supported prototypes, reducing time, material use, and cost while improving the likelihood that the final nozzle meets task-specific performance targets.

We therefore propose and study *task-specific multi-outlet nozzle* designs combining two elements: (i) a comparison of two archetypal multi-outlet geometries—a radial 90° splitter and a branched Y-splitter—across outlet counts; and (ii) evaluation against task-specific objectives that reflect practical priorities (high-throughput supports, precision patterning, and cell-safety-constrained extrusion). We focus on the mapping of geometry to application through quantitative metrics, thus moving beyond one-size-fits-all optimization. We build a reproducible CFD pipeline of 2- and 4-outlet nozzles with shear-thinning, power-law rheology representative of common bioinks (e.g., 10% gelatin methacrylate or GelMA, 2% methacrylated hyaluronic acid or MeHA, and 8% alginate) under pneumatic actuation (65 mbar to 105 mbar) (21). We compute spatial fields (static pressure, velocity magnitude, and WSS) and derive scalar metrics that capture performance: outlet flow imbalance, total pressure drop, and pressure-normalised throughput, defined as the mean outlet velocity divided by the inlet–outlet pressure drop (22). We report field maps, outlet symmetry, efficiency comparisons, and multi-metric trade-offs.

This work makes four specific contributions:

1. A *task-specific* evaluation framework for multi-outlet nozzles that formalises objectives for throughput, precision, and cell safety.
2. A comparative CFD study of two archetypes (90° radial and Y-split) at two outlet counts (2 and 4), showing how outlet number interacts with geometry to shape pressure loss, velocity, WSS, and flow balance.
3. Introduction and use of practical, decision-making metrics (outlet flow imbalance and pressure-normalised throughput, 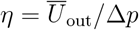.
4. Design guidance for practitioners: 90° 2-outlet nozzles as safe, balanced defaults for cell-laden printing; Y-split 2-outlet for fast, acellular deposition; and cautions for 4-outlet operation unless active balancing or control is available.

## 2 Methods

### 2.1 Methodology Overview

We assessed multi–outlet nozzle performance using a reproducible, simulation–driven workflow. The analysis considers three task classes: throughput-oriented printing (supports, acellular gels), cell-safety-constrained printing (cell-laden hydrogels), and precision deposition. We evaluate two archetypal geometries—a Y-split and a 90° radial splitter—at outlet counts of *N* = 2 and *N* = 4, using dimensions selected to match the constraints of the target bioprinter. Three representative bioinks (8% alginate, 2% MeHA, 10% GelMA) under 65 mbar, 85 mbar and 105 mbar pneumatic actuation. CAD geometries are imported into ANSYS Fluent, locally refined meshes are generated at junctions and bends, and steady laminar simulations are run using appropriate rheological models (power-law or Newtonian). From each simulation we extract outlet flow rates, mean outlet velocities, mean and peak WSS, the inlet–outlet pressure drop, and pressure-normalised throughput 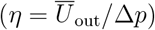. These scalar metrics are complemented by spatial field maps of pressure, velocity, and WSS. Finally, a qualitative assessment of the trade–offs of the resulting (speed, symmetry, shear, energetic cost) recommends task–specific geometries. Fig. 2 gives the general overview of our methodological computational framework, which is detailed in the following.

**Figure 1:**
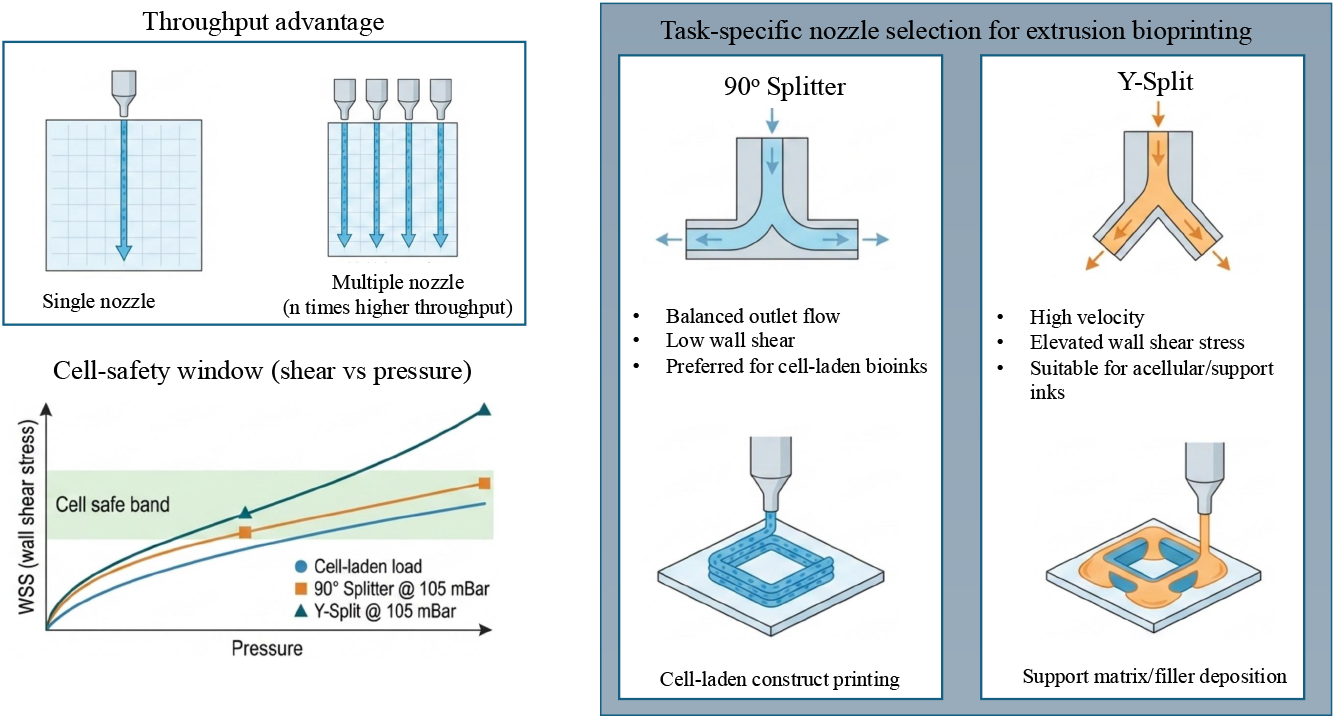
Overview of the extrusion bioprinting workflow, from CAD design to construct application. The process progresses through design, material preparation, printing, and post-print maturation before final assembly or implantation. This study focuses on the printing stage, where nozzle geometry dictates flow behaviour and cell stress.

**Figure 2:**
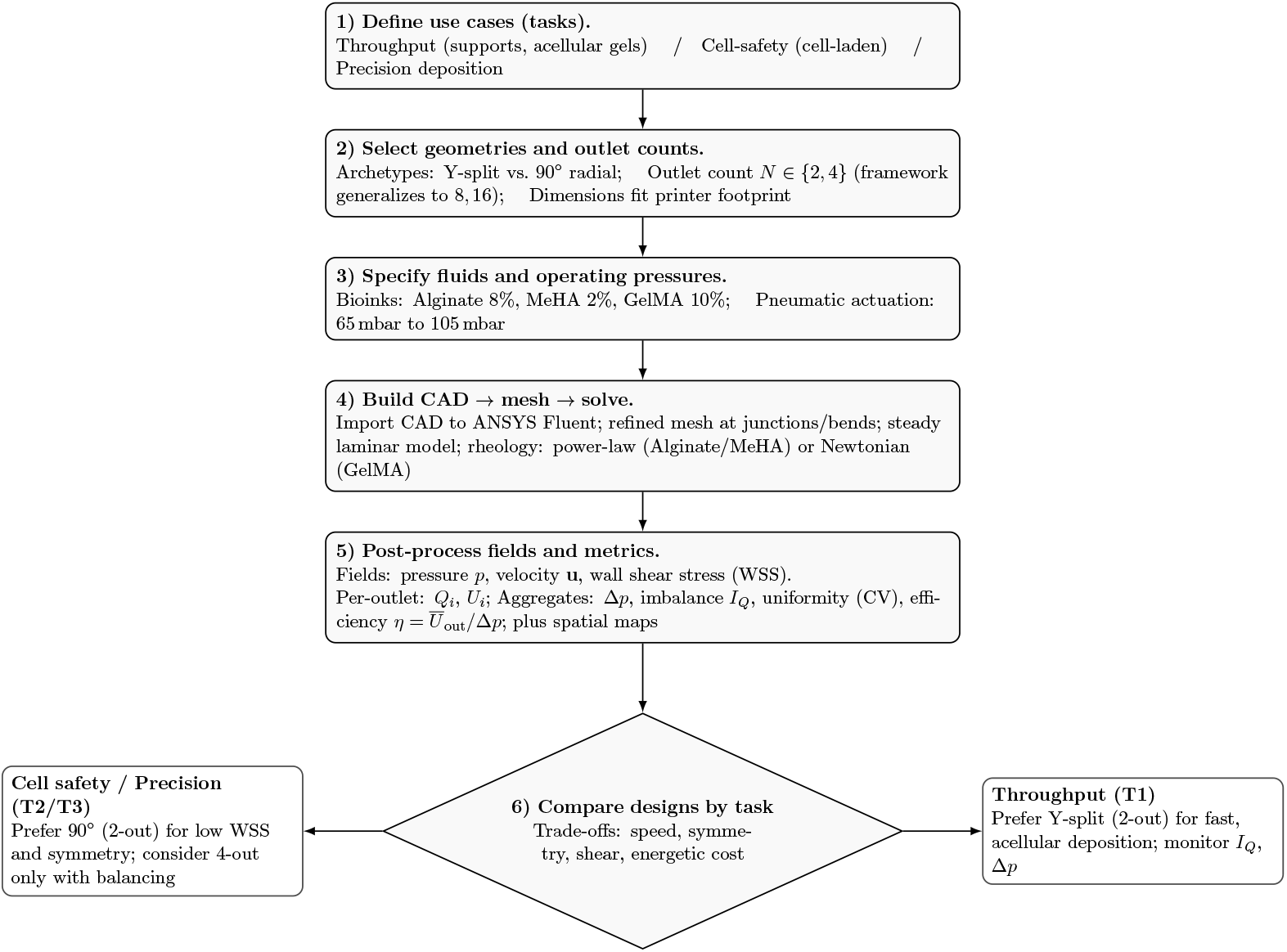
CFD-based workflow for task-specific evaluation of multi-outlet nozzles. Steps 1–5 define scenarios, solve the steady laminar problem with appropriate rheology, and extract comparable metrics. Step 6 maps results to task-oriented recommendations.

#### 2.1.1 Variables and Scenario Definition

To keep the study focused and interpretable, variables were grouped into *design, fluid*, and *operating* sets. Table 1 summarizes the parameter space explored in this study and makes explicit what was varied versus held fixed to enable fair comparisons across nozzle designs. On the design side, we varied only the outlet count (*N* = 2 or 4) and the branching archetype (Y–split versus 90° radial), while keeping interface-critical dimensions (outlet diameter, spacing, and needle length) constant to match the target printer hardware. Geometric details such as curvature and bifurcation angles were set by the chosen archetype, and the overall footprint was constrained by practical integration and sterilisation requirements. On the fluid and operating sides, we varied the bioink formulation and inlet pressure (65 mbar to 105 mbar gauge) while fixing outlet boundary conditions (0 mbar gauge), wall no-slip, and isothermal operation. This controlled scope isolates the influence of branching geometry and outlet count on flow balance and shear exposure. Table 3 specifies the bioink scenarios used across all nozzle configurations. Two inks (8% alginate and 2% MeHA) were modeled as shear-thinning power-law fluids with representative (*K, n*) parameters, whereas 10% GelMA was approximated as Newtonian with a constant viscosity (*μ* ≈0.46 Pa·s) under the simulated conditions. Density values were assigned per ink based on literature-informed assumptions to reflect modest formulation-dependent differences in inertia. These scenarios enable comparison of geometry effects across rheological regimes. We sampled 65 mbar, 85 mbar and 105 mbar to cover gentle cell-sparing operation (lower bound), typical lab settings (mid), and faster deposition regimes (upper bound), while remaining within the printer’s pressure capability and consistent with reported low-pressure extrusion ranges (50 mbar to 100 mbar) (21). Holding geometry and fluid properties fixed while varying pressure isolates each design’s hydraulic response, conversion of pressure into throughput, the resulting WSS, and sensitivity of outlet balance. This variation mirrors practical trade–offs between preserving cell viability (lower shear exposure) and increasing fabrication speed, and is consistent with CFD analyses showing that WSS increases with pressure under power-law rheology (22; 10).

**Table 1:** Study variables and scope. “Varied” indicates factors explored in this work; others were held fixed within practical constraints of the target printer and manufacturing route.

| Category | Variable | Scope in this study |
| --- | --- | --- |
| Design | Outlet count $N$ | Varied: $N = 2, 4$ (framework generalizes to 8, 16) |
|  | Branching geometry | Varied: Y-split vs. 90° radial |
|  | Outlet diameter, spacing, needle length | Fixed to fit printer hardware; consistent across cases |
|  | Curvature/taper, bifurcation angle | Set per archetype; smooth transitions for 90° design |
|  | Body thickness/footprint | Constrained by printer carriage and sterilisable envelope |
| Fluid | Bioink type | Varied: 8% alginate, 2% MeHA (power-law), 10% GelMA (Newtonian) |
|  | Density | Fixed per ink (Table 3) |
| | Rheology parameters | Fixed per ink (power-law $K, n$ or constant $\mu$) |
| Operating | Inlet pressure | Varied: 65 mbar, 85 mbar and 105 mbar (gauge) |
|  | Outlet condition | Fixed: 0 mbar (gauge), no-slip walls, isothermal |

### 2.2 Evaluation framework

Each simulation corresponds to a single nozzle configuration defined by its geometry, bioink, and inlet pressure. Geometry is varied only in outlet count (*N* = 2 or 4) and branching layout (Y–split or 90° split); all other dimensions are kept identical across designs. Operating conditions are set by the imposed inlet pressure and the assigned bioink rheology.

For each configuration, steady-state CFD simulations are used to compute internal velocity, pressure, and wall shear stress fields. From these fields, outlet flow rates, pressure drop, and mean and peak wall shear stress are extracted and used to compare nozzle behaviour.

#### Bioink rheology

The rheological behaviour of shear-thinning bioinks was modelled using a power-law (Ostwald–de Waele) formulation:

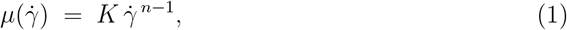

where *K* is the consistency index and *n* (*<* 1 for shear–thinning materials) controls how strongly viscosity falls with shear rate. Each bioink has its own parameters (*K, n*)—for instance, 10% GelMA is nearly Newtonian, whereas 8% alginate and 2% MeHA are strongly shear–thinning. For a given configuration *x*, the CFD solver computes velocity **u**(**r**), pressure *p*(**r**), shear rate 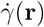, and wall shear stress 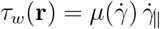 along the nozzle walls.

#### 2.2.1 Metrics per–outlet

In fluid mechanics, quantities such as velocity are not uniform across a surface and vary spatially due to boundary effects. For example, flow is typically faster near the centre of an outlet and slower near the walls due to viscous drag. To interpret the CFD results in physically meaningful terms, outlet-level metrics were derived from surface-integrated quantities computed directly from the velocity field.

The primary outlet metric is the *volumetric flow rate Q*_*i*_, which determines the printing throughput, i.e. the amount of material discharged per unit time from outlet *i ∈* 𝒪. It was defined as

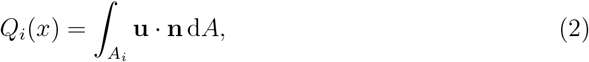

where **u** is the velocity field, **n** is the unit normal vector to the outlet cross-section, and *A*_*i*_ is the outlet area. This definition corresponds directly to the surface flux computed by the CFD solver.

A corresponding *mean outlet velocity* was derived consistently from the volumetric flow rate as

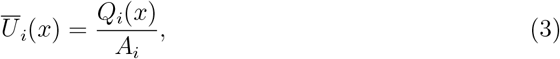

where 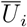 is the area-averaged normal velocity at outlet *i*. Because all outlet areas are identical, imbalance computed from *Q*_*i*_ is equivalent to imbalance computed from 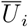.

Wall shear stress was not treated as a per-outlet metric. Instead, mean and peak wall shear stress were evaluated at the nozzle level as area-based quantities over the internal wetted wall surface, as defined in Section 2.2.2.

#### 2.2.2 Aggregate performance metrics

While per-outlet metrics reveal local behaviour, nozzle design decisions require system-level performance indicators that describe flow balance, hydraulic cost, and shear exposure within the nozzle.

To quantify how evenly flow is distributed across all outlets, the flow uniformity metric *ϕ*_*Q*_(*x*) was defined as

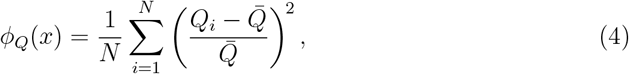

where *Q*_*i*_ is the volumetric flow rate at outlet *i*, 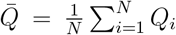 is the mean outlet flow rate, and *N* is the total number of outlets. This metric represents the normalised variance of outlet flow rates and provides a global measure of flow balance, with values approaching zero indicating near-uniform splitting.

Because averaged measures can mask severe deviations at individual outlets, a complementary worst-case metric was also defined. The flow imbalance *I*_*Q*_(*x*) was computed as

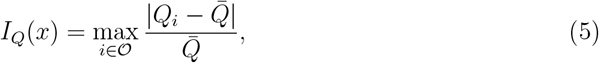

where 𝒪 = {1,..., *N*} denotes the set of outlets. This metric identifies the most over- or under-fed outlet relative to the mean flow rate.

The hydraulic cost of each design was characterised using the pressure drop

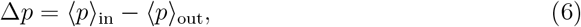

where ⟨ · ⟩denotes area-weighted averaging over the inlet and outlet planes. All pressures are reported as gauge pressures, with the outlet boundary prescribed at zero gauge pressure. Local regions of negative *gauge* pressure may appear in contour plots due to the chosen reference level (zero gauge at the outlets) and local acceleration effects. Design comparisons therefore use the pressure-drop metric Δ*p*, rather than interpreting local gauge minima as an outlet condition.

In addition to flow and pressure metrics, shear exposure along the nozzle walls was quantified to characterise the mechanical environment experienced by the bioink. The mean wall shear stress was defined as an area-weighted surface average over the internal wetted wall region,

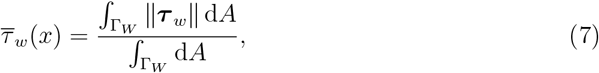

where ∥***τ*** _*w*_∥ is the wall shear stress magnitude and Г_*W*_ denotes the internal wall surface of the splitting block. The resulting value summarises the typical shear level within the nozzle.

Because localised shear peaks may occur even when the mean remains moderate, the peak wall shear stress was also evaluated as

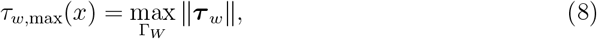

which identifies the maximum shear stress magnitude occurring anywhere on the same internal wall region Г_*W*_ and captures worst-case shear exposure.

Finally, pressure-normalised throughput was assessed using the efficiency index

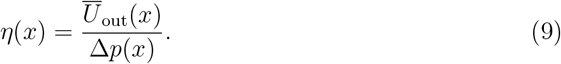

where 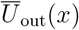 is the mean outlet velocity averaged across all outlets and Δ*p*(*x*) is the inlet–outlet pressure drop. This metric provides a compact measure of how effectively applied pressure is converted into outlet throughput.

#### 2.2.3 Task–specific performance criteria

Different bioprinting tasks place emphasis on different aspects of nozzle performance. High–throughput applications favour higher outlet velocities to increase deposition rate, whereas cell–laden printing prioritises reduced wall shear stress to limit shear–induced damage. Precision deposition, by contrast, requires highly uniform outlet flow together with stable, pressure–driven operation.

Rather than combining these competing requirements into a formal optimisation problem, nozzle designs were evaluated against task–specific performance criteria using directly computed CFD metrics. A throughput–oriented assessment focused on mean outlet velocity, interpreted alongside associated increases in wall shear stress and outlet flow imbalance. A cell–safety assessment emphasised both mean and peak wall shear stress as indicators of typical and worst–case shear exposure, with secondary consideration of flow uniformity. Precision–oriented assessment prioritised low outlet imbalance and moderate pressure drop while maintaining sufficient outlet velocity for stable extrusion.

Because practical printing conditions vary across bioink formulations and applied pressures, all designs were assessed consistently across the full set of simulated operating cases. Performance trends were therefore interpreted based on how robustly each geometry maintained acceptable velocity, shear, and flow balance across materials and pressure levels, rather than on behaviour under a single nominal condition.

To ensure biological and mechanical relevance, interpretation of the CFD results was guided by established feasibility limits reported in the literature. In particular, experimental studies have shown that high cell viability during extrusion is associated with limiting local wall shear stress and reducing the duration of cell exposure to high–shear regions (23). Similarly, practical printer constraints—including maximum allowable inlet pressure, manufacturable channel dimensions, and acceptable outlet flow imbalance as outlet count increases—were used to contextualise the results and identify designs likely to remain viable when scaled.

Together, these task–specific criteria define a realistic evaluation framework for comparing nozzle geometries. The analysis links CFD–resolved velocity, pressure, and wall shear stress fields directly to practical printing objectives without introducing a formal optimisation model or weighted objective function.

### 2.3 Designs

The nozzle geometry governs how flow divides and how shear develops during extrusion. To isolate geometric effects, four configurations were studied: two–outlet Y–split, two–outlet 90° split, four–outlet Y–split, and four–outlet 90° split. All models shared a common inlet channel (*inlet length L*_in_ = 8.50 mm) and identical outlet diameters (*outlet diameter D*_out_ = 1.35 mm) elected to match a commonly used extrusion scale for hydrogel bioprinting. Channel dimensions were chosen to match the resolution limits of the 3D-printed moulds (printable channels ≳1.2 mm, with walls thick enough to avoid leakage under pneumatic loading) and to remain within the bioprinter’s pressure limit (≤ 200 mbar). Complete geometry specifications for all nozzle designs are reported in Tables 7–10.

To ensure reproducibility while enabling meaningful comparison, each geometry was defined by a small set of *template parameters*:

- **Outlet count** *N ∈ {*2, 4*}*.
- **Branching archetype:** Y–split (acute bifurcation) vs. orthogonal (90°) split.
- **Junction geometry:** For Y–split designs, the bifurcation angles θ_Y_ and all other branching parameters are reported in the corresponding geometry tables. For 90° split designs, flow redirection was implemented using tangent–arc bends with a single, fixed bend radius *R*_*c*_ defined per configuration.
- **Curvature control:** the internal bend radius *R*_*c*_ and the straight sections before and after each turn were sized to minimise recirculation.
- **Transition shaping:** no taper within the split-block (*α*=0); tapering is applied only in the shared downstream needle geometry.

Dimensions were set to remain within the printhead’s available space and to preserve left-right symmetry, so that the flow paths on both sides had the same length and hydraulic resistance. Table 2 lists only geometry parameters common to all designs. All branching-specific dimensions are reported separately.

**Table 2:** Geometric parameters shared by all nozzle configurations. Exact split-block geometries are specified per nozzle in Tables 6–9.

| Item | Symbol / Value | Rationale |
| --- | --- | --- |
| Inlet straight length | $L_{\text{in}} = 8.50 \text{ mm}$ | Provides a stable, fully developed inlet flow before the first junction. |
| Channel diameter (split-block) | $D = 1.35 \text{ mm}$ | Constant diameter throughout the split-block to isolate geometric effects and ensure fair comparison across designs. |
| Outlet diameter (at split-block exit) | $D_{\text{out}} = 1.35 \text{ mm}$ | Fixed outlet size used across all configurations; shared needle geometry is defined separately (Table 10). |
| Minimum printable diameter | $\geq 1.2 \text{ mm}$ | Ensures manufacturability and reduces clogging risk for hydrogel extrusion. |
| Outlet count | $N = 2 \text{ or } 4$ | Baseline and first scalable multi-outlet configurations studied. |
| Internal taper (split-block) | $\alpha = 0$ | No taper within the split-block; all channels maintain constant diameter. Tapering is applied only in the downstream needle section. |
| Build planarity | single build plane | Simplifies fabrication, cleaning, and meshing while preserving symmetry. |

#### 2.3.1 Two–outlet designs

The two–outlet geometries use a single bifurcation from the inlet to two identical outlets and serve as the baseline for assessing geometric effects. Each layout is mirror-symmetric, with the same path length and cross-section on both sides, so the ideal outcome is a 50:50 flow split. *Two–outlet Y–split:* The junction uses an acute bifurcation (*θ*_Y_) to turn the flow gradually and reduce junction losses. Short tangent sections were added after the split to let the flow stabilise before entering the final taper and outlet. This design prioritises smoother streamlines and higher throughput at a given pressure, but requires a longer overall length. *Two–outlet 90*° *split:* Flow is redirected into perpendicular branches via a single–radius fillet (*R*_*c*_) rather than a sharp right angle. The orthogonal layout is more compact and straightforward to fabricate and scale. The tighter turning can raise local WSS, but the symmetry and short path lengths favour balanced outlet flows and moderate pressure drops.

#### 2.3.2 Four–outlet designs

The four–outlet variants add a second, symmetric split to each branch (inlet → 2 → 4). The second junction duplicates the first (same design and shaping) to preserve path–length equality from inlet to all outlet. This layout tests the first stage of scaling: whether any small imbalance or high-shear region created at the first split becomes larger or smaller after the second split. *Four–outlet Y–split:* The four-outlet Y-split uses a two-stage angle strategy: a wider primary split followed by a narrower secondary split to keep flow redirection gentle at both junctions. This approach produces a smoother, more gradual branching layout, though at the cost of a longer overall path. *Four–outlet 90*° *split:* In the four–outlet 90° split, each branch undergoes a second orthogonal split using the same bend radius *R*_*c*_ and tangent lengths as the first tier, preserving geometric consistency across both stages.

### 2.4 Bioink Models

#### 2.4.1 Density and Viscosity

Density enters the inertial term and is retained for Reynolds-number estimation; in these viscous, low-Reynolds-number nozzle flows its effect on pressure drop and WSS is secondary to viscosity. Differences among the selected bioinks are small relative to viscosity variations but remain relevant for force transmission and for *a posteriori* Reynolds number checks. Accordingly, representative density values were assigned consistent with water–based hydrogels: 1050 kg*/*m^3^ for 8% alginate, 1030 kg*/*m^3^ for 10% GelMA, and 1000 kg*/*m^3^ for 2% MeHA.

The alginate density value follows experimentally measured densities for extrusion-relevant alginate bioinks (5–15 wt%), which are reported to lie near 1.05 g cm^−3^ (24). The GelMA density value is consistent with experimentally measured densities for 10 wt% gelatin hydrogels (approximately 1.025 g cm^−3^), from which GelMA is derived (25). For MeHA, density was taken equal to that of water, reflecting its dilute, water-dominated formulation (approximately 98 wt% water) and consistent with literature treatments of HA/MeHA bioinks in which density deviations are negligible and not treated as independent material parameters (26). These values are treated as modelling assumptions rather than tuned material parameters, avoiding artificial inflation of density variability while maintaining consistency across scenarios in which viscosity dominates the flow behaviour.

Across all cases, Reynolds numbers based on the outlet diameter *D*_out_, the mean outlet velocity, and the effective viscosity were 𝒪 (1) or lower. The flow is therefore viscous-dominated and laminar, with wall shear stress (WSS) governed primarily by local velocity gradients rather than inertial effects. A 5% variation in density alters Reynolds numbers proportionally but does not change the flow regime or the relative ranking of nozzle designs.

To reflect extrusion-relevant rheology, 8% (w/v) alginate was modelled as a shear-thinning, power-law fluid. Alginate-based bioinks are widely reported to exhibit decreasing apparent viscosity with increasing shear rate under extrusion-scale conditions, and are commonly described using power-law or Herschel–Bulkley models over shear rates relevant to bioprinting (27).

For MeHA (2% w/v), a power-law representation was adopted over the nozzle-relevant shear-rate window. HA- and MeHA-based bioinks are widely reported to exhibit pronounced shear-thinning behaviour under extrusion conditions, with viscosity–shear-rate flow curves used to define printing pressures and operating regimes. Over shear rates relevant to extrusion bioprinting, simple non-Newtonian constitutive models, such as power-law or Herschel–Bulkley formulations, are commonly used to represent the measured rheology (26).

For both alginate and MeHA, the power-law parameters (*K, n*) were selected to represent reported steady-shear behaviour over shear rates relevant to nozzle extrusion and were held constant for each bioink across all simulations. In contrast, 10% (w/v) GelMA at 37 °C was modelled as a near-Newtonian fluid using a representative apparent viscosity (*µ*_*G*_ = 0.46 Pa s). This value is not treated as a unique material constant, rather, it is consistent with viscosities reported for 10% GelMA at near-physiological temperature over extrusion-relevant shear rates, where shear dependence is markedly reduced (28, Sec. 3.1, Fig. 3), and lies within the sub-Pa s range commonly reported for GelMA solutions at physiological temperature in steady-shear and extrusion contexts (29). The viscosity *µ*_*G*_ is therefore adopted here as a representative modelling input to enable controlled comparison of nozzle geometries under fixed rheological conditions.

**Figure 3:**
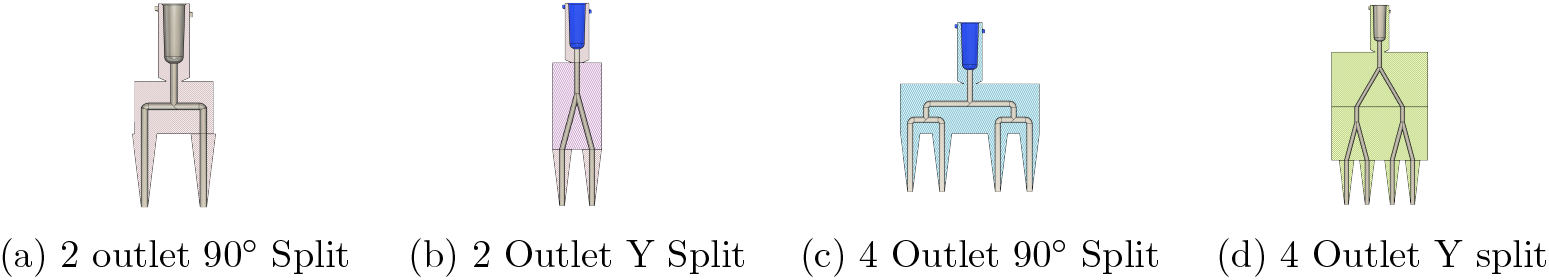
CAD renderings of the four multi-outlet nozzle configurations evaluated in this study: (a) 2-outlet 90° split, (b) 2-outlet Y-split, (c) 4-outlet 90° split, and (d) 4-outlet Y-split. All geometries share the same inlet and outlet diameters and were exported from the same STEP models used for meshing and CFD.

##### Shear-thinning bioinks (alginate, MeHA)

Sodium alginate (8%) and MeHA (2%) were represented as generalized Newtonian shear-thinning fluids whose viscosity followed a power-law dependence on shear rate

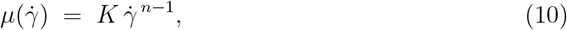

where *μ* is the apparent viscosity, 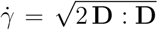 is the scalar shear rate computed from the rate–of–strain tensor **D**, *K* is the consistency index (units: Pa s^*n*^), and *n* ∈ (0, 1) is the flow index for shear–thinning fluids. Power–law parameters (*K, n*) were obtained by fitting literature–reported steady–shear viscosity data for printable alginate and MeHA bioink formulations over the shear–rate range relevant to extrusion ( ≈ 10^1^–10^4^ s^−1^), following established bioprinting rheology practice (30; 24; 26; 31; 32). The resulting parameters are treated as representative modelling inputs rather than as calibrated material constants. To maintain numerical stability and avoid unphysical viscosity values, a *regularized* form was used:

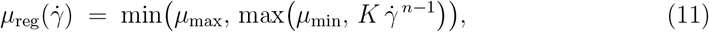

with bounds (*μ*_min_, *μ*_max_) chosen to prevent singular behaviour at low shear and unrealistically low viscosities at high shear. The shear rates generated in the nozzle (order 10^1^–10^4^ s^−1^) fall within the range over which the rheological parameters were fitted (29). Unless stated otherwise, we used *µ*_min_ = 0.001 Pa s and *µ*_max_ = 10 Pa s (Table 3) to prevent singular behaviour at low shear and unrealistically low viscosities at high shear.

**Table 3:** Parameter values are representative power–law fits to literature rheology data over extrusion–relevant shear rates and are used for comparative CFD analysis rather than calibrated material properties.

| Bioink | Model | Parameters | Bounds | Density [kg/m<sup>3</sup>] |
| --- | --- | --- | --- | --- |
| Alginate 8% | Power-law | $K = 56.9 \text{ Pa} \cdot \text{s}^n$, $n = 0.322$ | $\mu_{\min} = 0.001 \text{ Pa} \cdot \text{s}$, $\mu_{\max} = 10 \text{ Pa} \cdot \text{s}$ | 1050 |
| MeHA 2% | Power-law | $K = 12.4 \text{ Pa} \cdot \text{s}^n$, $n = 0.55$ | $\mu_{\min} = 0.001 \text{ Pa} \cdot \text{s}$, $\mu_{\max} = 10 \text{ Pa} \cdot \text{s}$ | 1000 |
| GelMA 10% | Newtonian | $\mu_G = 0.46 \text{ Pa} \cdot \text{s}$ | – | 1030 |

##### Near-Newtonian bioink (GelMA)

GelMA (10%) was treated as near-Newtonian over the shear-rate range relevant to the present nozzle flows (29). Accordingly, a constant apparent viscosity was assumed,

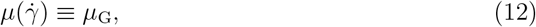

where *μ*_G_ is selected as a representative steady–shear viscosity within the mid–range of nozzle-relevant shear rates. This modelling choice reflects the comparatively weak shear–rate dependence reported for GelMA at similar concentrations in extrusion and bioprinting studies, where steady–shear viscosities are commonly used instead of full non– Newtonian fits. This approximation reduces model complexity and provides a controlled baseline for comparing geometries under fixed rheological conditions.

Because power–law models are typically characterised over a finite shear–rate window, a regularised viscosity formulation was employed to prevent the model from producing unrealistically large or small viscosities outside that range. The bounds (*μ*_min_, *μ*_max_) in (11) were selected to confine the apparent viscosity to values commonly reported for hydrogel extrusion, spanning both low– and high–shear conditions (33). These limits do not represent yield behaviour; instead, they ensure stable behaviour when local shear rates fall outside the region over which power–law behaviour is usually reported.

For GelMA (10%), a near–Newtonian response was assumed under the operating conditions considered here, and a single representative viscosity *µ*_G_ was used. This value corresponds to the mid–range of shear rates expected during extrusion and reflects the comparatively weak shear dependence reported for GelMA at similar concentrations. Each bioink was assigned a constant density *ρ*.

To confirm the appropriateness of the laminar flow assumption, Reynolds numbers were evaluated using the CFD-resolved mean outlet velocity *U* and outlet diameter *D*,

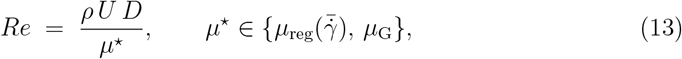

where *µ* ^⋆^ denotes the effective viscosity associated with each material model.

Table 3 summarises the rheology models and parameters adopted for the CFD simulations. The numerical values of (*K, n*) and *μ*_G_ are not intended to represent unique material characterisations, but to provide consistent rheological conditions for comparing nozzle geometries. Accordingly, identical parameter choices were applied across all designs, and their influence on predicted flow behaviour was examined through sensitivity analysis.

### 2.5 Performance Metrics

Performance was assessed using the same four metrics defined in §2.2: (i) wall shear measures (mean and peak WSS) as a proxy for cell safety, (ii) per-outlet and total volumetric flow for throughput and outlet balance, (iii) pressure field and overall pressure drop for hydraulic cost and printer constraints, and (iv) outlet velocity as a complementary indicator of deposition behaviour. Metrics are reported and interpreted using those prior definitions without redefinition, and are discussed separately rather than collapsed into a single objective to reflect the inherent trade-offs between throughput, balance, and shear exposure.

#### Wall shear

Mean WSS 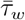 (area-averaged over the split-block walls) and peak WSS *τ*_*w*,max_ were used to capture typical shear exposure and local hotspots at junctions and turns. Wall shear stress was evaluated directly from the CFD solution and is used as a comparative indicator of mechanical loading relevant to cell-laden extrusion, consistent with prior studies on shear-mediated cell damage. (34; 35; 36).

#### Flow rate and balance

Outlet flow rates *Q*_*i*_ quantify throughput and provide the basis for the flow uniformity and imbalance measures in §2.2. Reporting both Σ_*i*_ *Q*_*i*_ and the outlet-to-outlet distribution isolates how geometry affects scalability and deposition uniformity in multi-outlet extrusion.

#### Pressure

Pressure drop Δ*p* between inlet and outlet planes was used as the primary energetic constraint, together with hydraulic power 𝒫_*h*_ = Δ*p* Σ_*i*_ *Q*_*i*_. Pressure contours were used only to localise dominant loss regions; local gauge extrema are shown for interpretation and are not treated as an outlet “suction” metric.

#### Velocity

Velocity fields and outlet-mean velocities (derived from *Q*_*i*_) were reported as complementary indicators of flow redistribution and potential deposition differences across outlets. For shear-thinning inks, velocity variations are coupled to local viscosity through 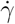-dependent rheology, linking outlet balance and shear hotspots to the same underlying flow features.

### 2.6 CFD simulations

Computational fluid dynamics (CFD) is widely used to resolve in-nozzle pressure, velocity, and wall shear stress fields in extrusion bioprinting, which are difficult to access experimentally due to the small scale and opacity of nozzle channels (37; 38; 32; 22; 39; 13; 40). Here, CFD provides a controlled basis for comparing multi-outlet nozzle geometries under identical operating conditions.

All simulations were performed in ANSYS Fluent 2025 R1 assuming steady, incompressible, laminar flow. Shear-thinning bioinks were represented using a power-law generalized Newtonian viscosity, while GelMA was treated using a constant-viscosity approximation (41). The governing equations are

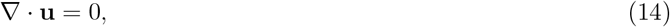

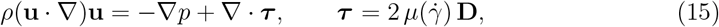

where **u** is the velocity, *p* the pressure, *ρ* the density, 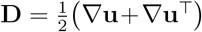 the rate-of-strain tensor, and 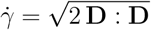 the scalar shear rate.

Across all cases, Reynolds numbers based on outlet diameter *D*, mean outlet velocity *U*, and an effective viscosity *µ*^⋆^ were 𝒪(1) or lower, supporting the laminar assumption. A pressure inlet was applied with *p*_in_ ∈ {65 mbar, 85 mbar, 105 mbar} (gauge) and all outlets were set to *p*_out_ = 0 mbar (gauge); walls were no-slip, and temperature and gravity effects were neglected.

Each geometry (Y-split and 90^°^ split) and outlet count (*N* = 2, 4) was meshed separately with local refinement at junctions and outlets and near-wall layers to resolve velocity gradients. A pressure-based solver with SIMPLE coupling was used. Gradients were computed by least-squares, and second-order spatial discretisation was applied for pressure and momentum. Apparent viscosity was updated from the local shear rate at each iteration following Fluent’s standard non-Newtonian implementation (41). For numerical stability at very low strain rates, viscosity limits were enforced for power-law inks. The upper bound (*µ*_max_ = 10 Pa s) was reached locally in near-stagnant regions for alginate and MeHA, with verification of the realised viscosity range reported in Supplementary S2.

Convergence was assessed by scaled residuals (*<* 10^−5^ for continuity and momentum), global mass imbalance (*<* 0.1%), and stabilisation of monitored area-averaged outlet flow rates. Simulations were advanced to approximately 10^3^ iterations to ensure steady fields. For representative cases, grid-refinement studies were performed until changes in area-averaged outlet velocity 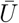 and mean wall shear stress 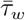 were below 1% and 2%, respectively, indicating mesh-independent results (42). Unless stated otherwise, identical boundary conditions, material parameters, and meshing strategy were applied across cases to enable direct geometry-to-geometry comparisons. All pressures are reported as gauge relative to the outlet boundary (0 mbar). Inlet pressures are specified in mbar to reflect printer settings, while pressure fields are reported in pascals as returned by the solver (1 mbar = 100 Pa).

## 3 Results

### 3.1 Spatial Field Analysis

Figure 4 contrasts the 2-outlet geometries for 8% alginate at 105 mbar. Relative to the 90^°^ split, the Y-split shows higher WSS and a more centralised high-velocity core, indicating a larger region of elevated shear within the splitter while sustaining higher branch velocities. Static pressure contours are reported as gauge relative to the outlet boundary (*p*_out_ = 0) and are included for qualitative interpretation only. Locally negative gauge values may therefore appear and are not used as a suction metric.

**Figure 4:**
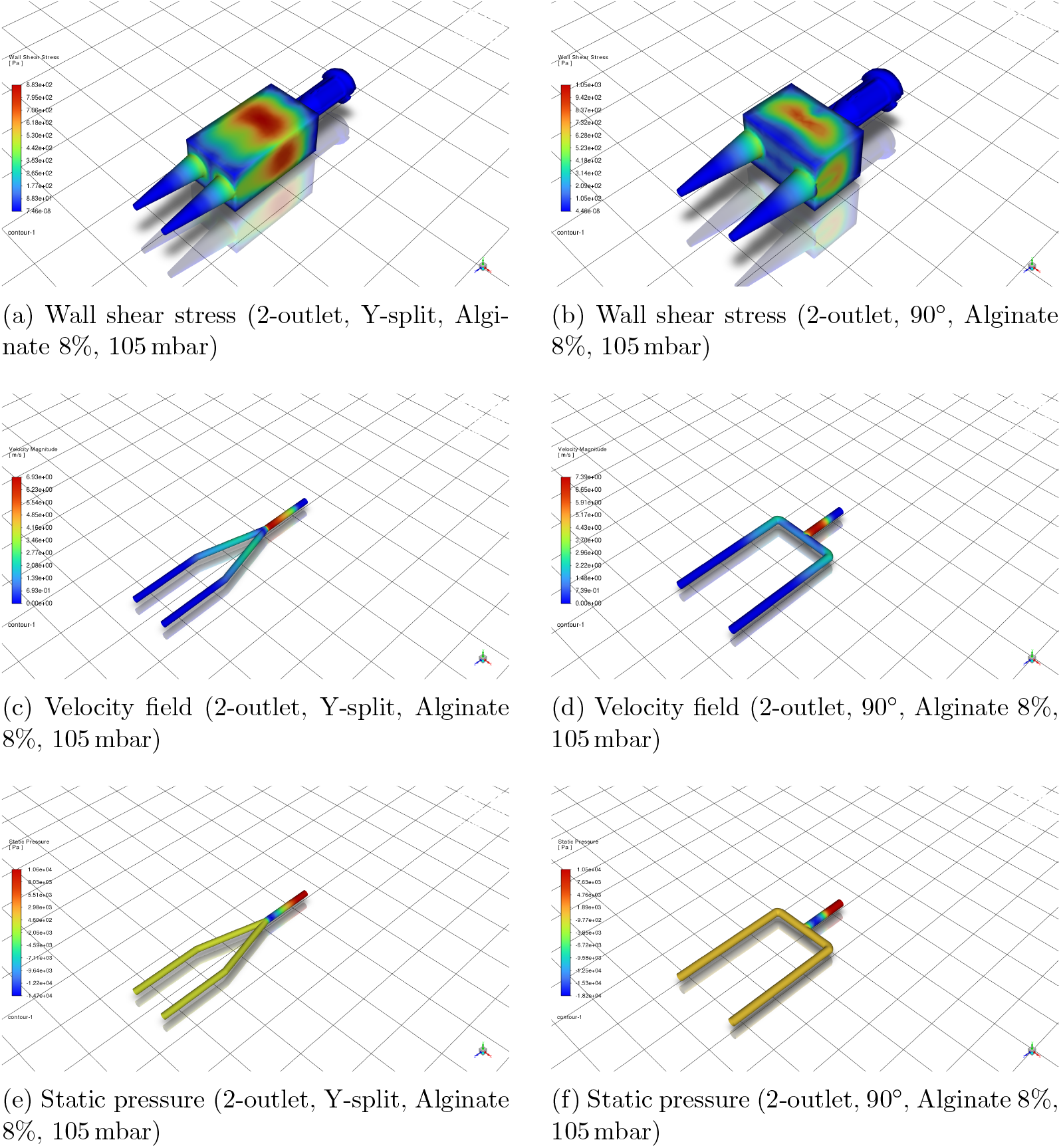
Comparison of 2-outlet nozzle designs (Y-split vs. 90^°^) across key spatial metrics for Alginate 8% at 105 mbar.

Figure 5 presents representative spatial fields for selected four-outlet cases. At 105 mbar, the four-outlet 90^°^ design exhibits a visually symmetric velocity field, whereas the four-outlet Y-split shows peak WSS localised at the primary junction, indicating that the dominant shear hotspot persists with increasing outlet count. At lower inlet pressure, the branched network becomes more susceptible to maldistribution, consistent with the outlet-imbalance trends (Fig. 7).

**Figure 5:**
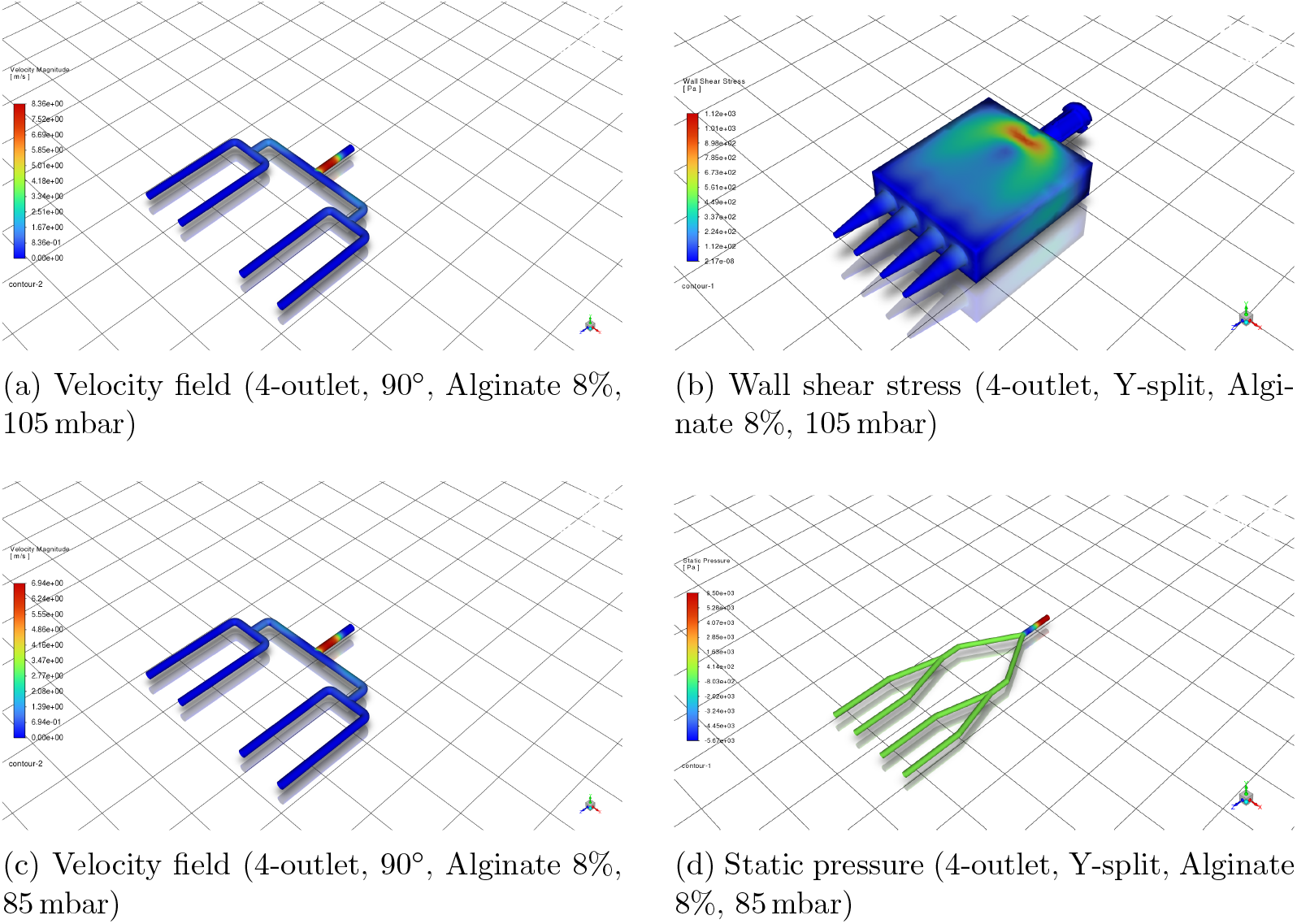
Spatial distribution of velocity, wall shear stress, and static pressure across selected nozzle designs and test cases.

Table 4 summarises the main findings across all simulated nozzle configurations. Flow balance was highest for the 2-outlet 90^°^ split design, with *I*_*Q*_ below 1% for shear-thinning bioinks (alginate and MeHA), while GelMA exhibited substantially higher imbalance. This configuration also exhibited the lowest wall shear stress (WSS), indicating suitability for cell-laden bioinks where shear-induced damage must be minimised (43). The 2-outlet Y-split geometry achieved higher outlet velocities, favouring throughput-driven applications (e.g., support-structure printing), but incurred higher WSS and moderate flow imbalance. Increasing the outlet count to four reduced flow symmetry in both designs. Y-split geometries exhibited the highest shear stress concentrations (frequently exceeding 150 Pa). They also exhibited lower pressure-normalised throughput compared with the corresponding 90^°^ designs. The 4-outlet 90^°^ design provided a more balanced compromise, maintaining lower WSS and improved symmetry than the 4-outlet Y-split, though remaining inferior to the 2-outlet configurations Overall, this analysis supports a task-oriented nozzle selection strategy: (i) Use 90^°^, 2-outlet for precision and cell safety, (ii) Use Y-split, 2-outlet for fast printing and bulk deposition, (iii) Avoid 4-outlet geometries for delicate bioinks unless active compensation or flow control is implemented. All the code and STEP files are uploaded to github, at https://github.com/Ceurda/bioprinter-nozzle-paper.

**Table 4:** Qualitative summary of performance trade-offs across nozzle configurations and metrics. Ratings reflect relative behavior from CFD results.

| Configuration | Flow Balance | WSS (Cell Safety) | Pressure loss localisation | Velocity | Efficiency ($\eta = \overline{U}_{out}/\Delta p$) |
| --- | --- | --- | --- | --- | --- |
| 2-Outlet, 90° Split | ✓✓✓ | ✓✓✓ | ✓✓✓ | ✓ | ✓✓✓ |
| 2-Outlet, Y-Split | ✓✓ | ✓ | ✓✓ | ✓✓✓ | ✓ |
| 4-Outlet, 90° Split | ✗ | ✓✓ | ✓✓ | ✓✓ | ✓✓ |
| 4-Outlet, Y-Split | ✗ | ✗ | ✓ | ✓✓✓ | ✗ |

### 3.2 Cross-condition comparison across bioinks and inlet pressures

In Figure 6a, static pressure contours indicate that pressure losses in Y-split geometries are primarily localised at the primary bifurcation, whereas in the 90^°^ designs losses are distributed across successive bends and straight sections. These differences reflect local acceleration, turning, and junction effects within the nozzle and are used here solely to diagnose loss localisation rather than absolute pressure magnitude. Quantitative comparison between designs is therefore based on Δ*p*, as defined in §2.2.

**Figure 6:**
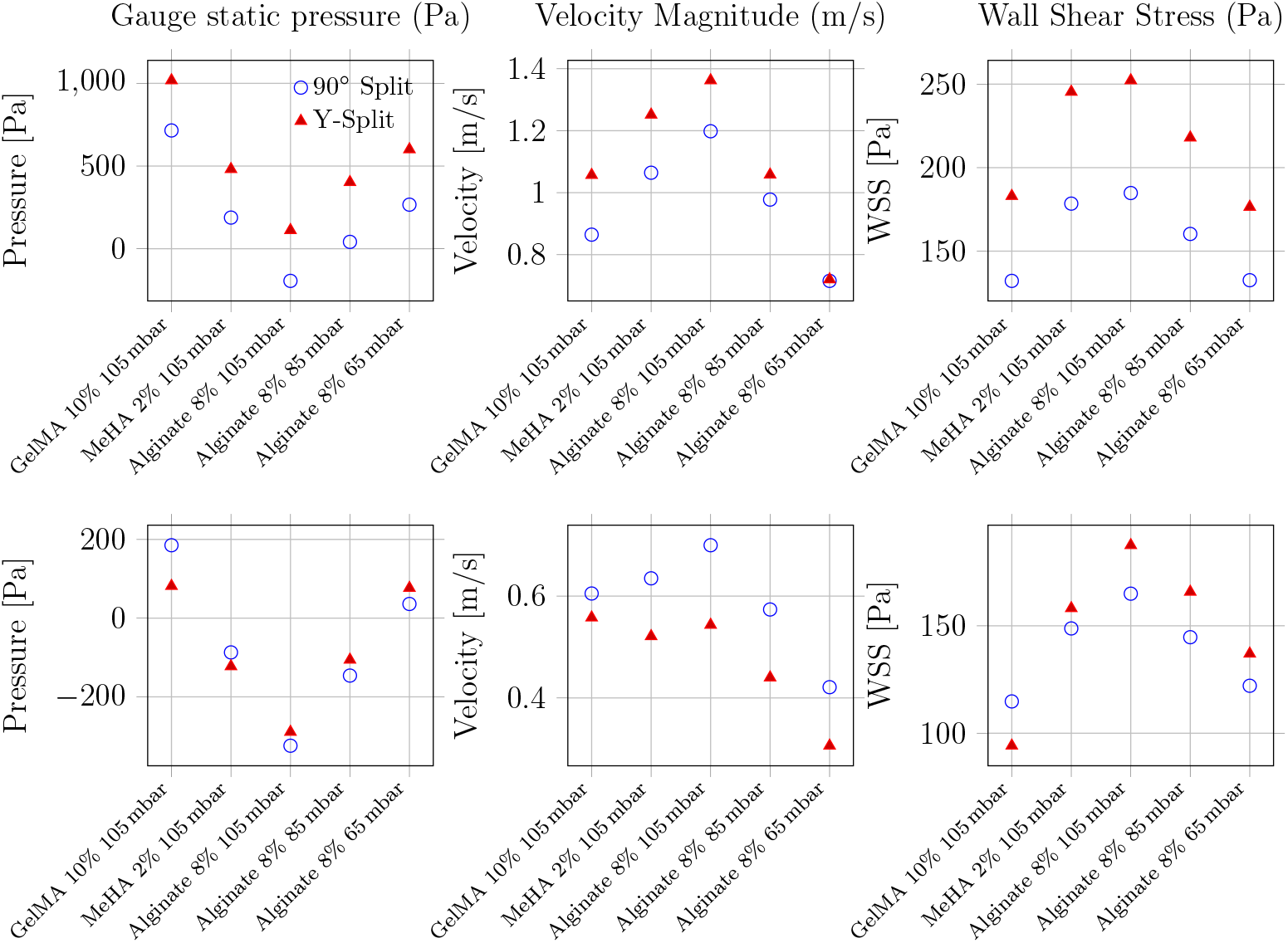
CFD-based performance comparison of Y-split and 90^°^ nozzles for two- and four-outlet configurations. Top row: 2 outlets. Bottom row: 4 outlets. Columns show static pressure, velocity magnitude, and wall shear stress. Static pressure is shown as gauge pressure relative to the outlet boundary (*p*_out_ = 0) and is included to visualise loss localisation rather than absolute pressure magnitude.

In terms of velocity, Y-split designs also produce consistently higher outlet speeds, particularly for lower-viscosity MeHA and alginate. At 105 mbar, Y-split nozzles exceed 1.3 m/s, compared with approximately 1.2 m/s in 90^°^ designs. This difference reflects reduced minor losses associated with smoother bifurcation in the Y-split geometry.

However, these benefits come at a cost: wall shear stress (WSS) is consistently higher in the Y-split configuration, particularly under high inlet pressure and low-viscosity conditions. Maximum WSS reaches approximately 250 Pa for 8% alginate at 105 mbar. Reported shear–viability thresholds in extrusion bioprinting vary widely with cell type, bioink formulation, and nozzle residence time. Accordingly, WSS is interpreted here as a comparative indicator of relative shear exposure rather than as a universal damage threshold (44). In this comparative context, the 90^°^ nozzle exhibits a more conservative shear profile across all tested cases, supporting its suitability for cell-laden bioinks.

Figure 6b extends the analysis to four-outlet configurations. In four-outlet configurations, both geometries exhibit reduced outlet velocity and increased sensitivity to pressure and bioink rheology. The static-pressure fields become more spatially uniform with increasing outlet count, indicating redistribution of hydraulic losses across additional branches. Yet, the Y-split design continues to exhibit slightly higher outlet velocities, mirroring trends from the two-outlet case.

The velocity advantage of the Y-split design diminishes in the four-outlet configuration. The performance gap between geometries narrows, particularly under lower-pressure conditions (e.g., 65 mbar with 8% Alginate), where both designs converge to similar velocity magnitudes ( 0.5–0.6 m/s). This trend suggests that increasing outlet count reduces the benefit of smoother Y-transitions, likely due to cumulative minor losses and symmetry degradation.

The WSS trends, however, persist: Y-split geometries still produce higher peak shear values, especially for high-pressure, low-viscosity inputs. The 90^°^ nozzle again demonstrates a more conservative shear profile, implying safer use for bioinks with embedded living cells. For example, GelMA at 105 mbar yields approximately 110 Pa in WSS for the 90^°^ layout versus 135 Pa in the Y-split.

### 3.3 Impact of different materials

Figure 7 quantifies outlet non-uniformity using the imbalance metric *I*_*Q*_ across bioinks and inlet pressures for the two- and four-outlet designs. For the two-outlet geometries (Fig. 7a–b), *I*_*Q*_ remains negligible for the shear-thinning bioinks (alginate and MeHA), staying below 1.5% across all tested inlet pressures. GelMA is the clear outlier: at 105 mbar, *I*_*Q*_ increases to 14.8% for the 90^°^ split (Fig. 7a) and 31.3% for the Y-split (Fig. 7b), indicating substantially higher sensitivity of the split to the GelMA operating regime.

**Figure 7:**
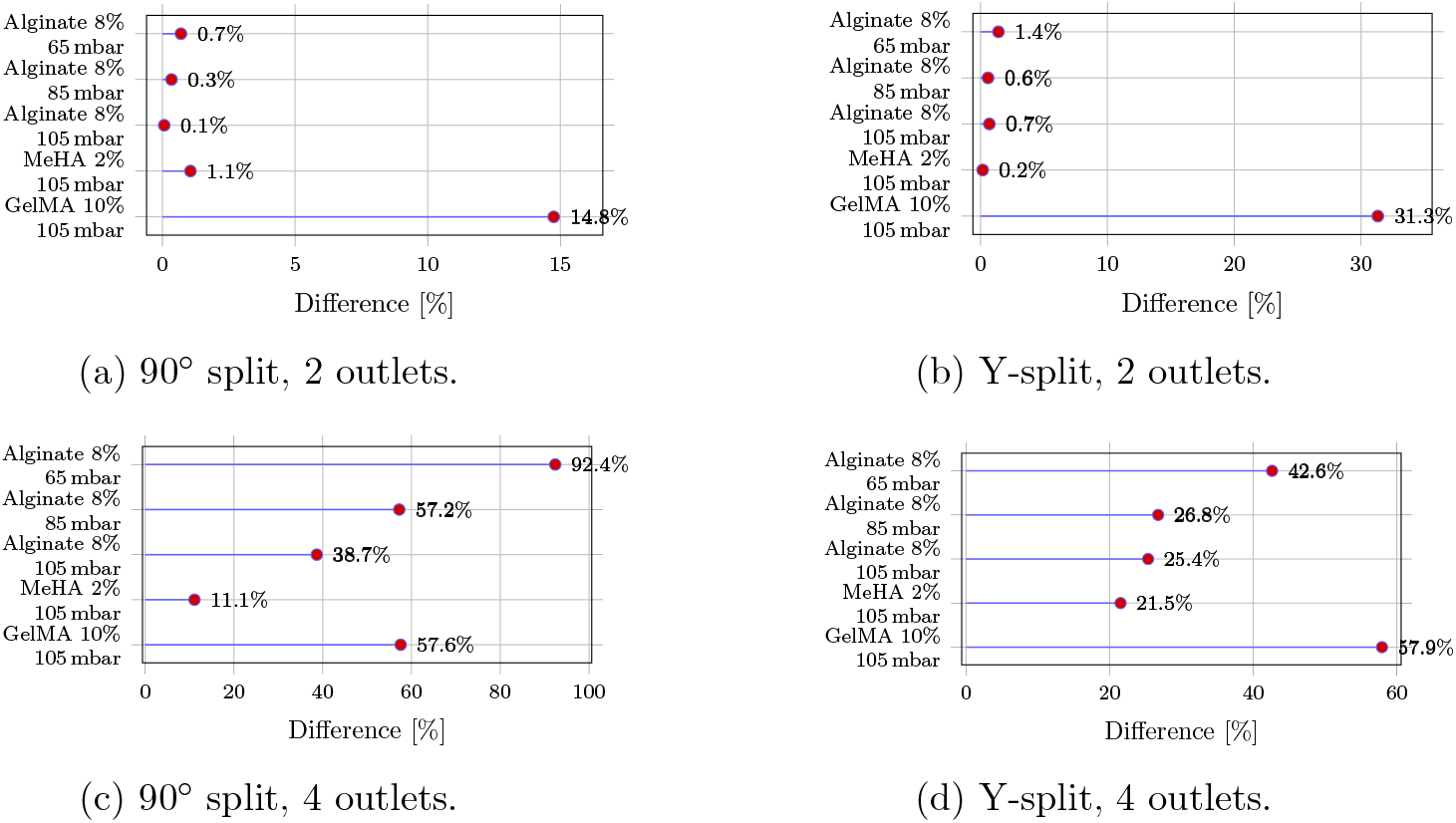
Outlet flow imbalance was quantified using the flow-rate imbalance metric *I*_*Q*_ defined in §2.2, computed from the volumetric flow rates *Q*_*i*_ on each outlet plane. Because all outlet areas are identical, the same trends are reflected in outlet mean velocities, but all imbalance values reported here are based on *Q*_*i*_ for consistency with the primary throughput metric. Figure 7 reports the flow-rate imbalance metric *I*_*Q*_ across all outlet counts and operating cases. Subfigures (a) and (b) compare the 2-outlet configurations, while (c) and (d) extend the comparison to 4-outlet designs.

In contrast, the four-outlet configurations (Fig. 7c–d) exhibit a pronounced loss of uniformity across all materials. Flow-rate imbalance exceeds 20% in all four-outlet cases and reaches 57.9% for GelMA in the four-outlet Y-split configuration. (Fig. 7d). The four-outlet 90^°^ manifold is additionally pressure-sensitive, with asymmetry worsening at lower inlet pressure and peaking at 92.4% for alginate at 65 mbar (Fig. 7c). Overall, Fig. 7 shows that two-outlet designs provide robust outlet uniformity for shear-thinning bioinks, whereas four-outlet splitting introduces strong sensitivity to rheology and inlet pressure that would likely require active compensation to maintain uniform deposition.

### 3.4 Integrated Performance Metrics and Trade-offs

Figure 8 compares pressure-normalised throughput *η* across all tested conditions for two-outlet nozzles. Across all cases, the 90^°^ design exhibits higher efficiency values than the Y-split counterpart. This indicates that although the Y-split achieves higher outlet velocities, it does so with a greater pressure cost (higher Δ*p*), reducing pressure-normalised throughput under fixed pneumatic actuation. The 90^°^ design maintains more favourable pressure-normalised throughput, indicating lower hydraulic cost under fixed pneumatic actuation. Cell viability considerations are assessed separately using mean and peak wall shear stress (45).

**Figure 8:**
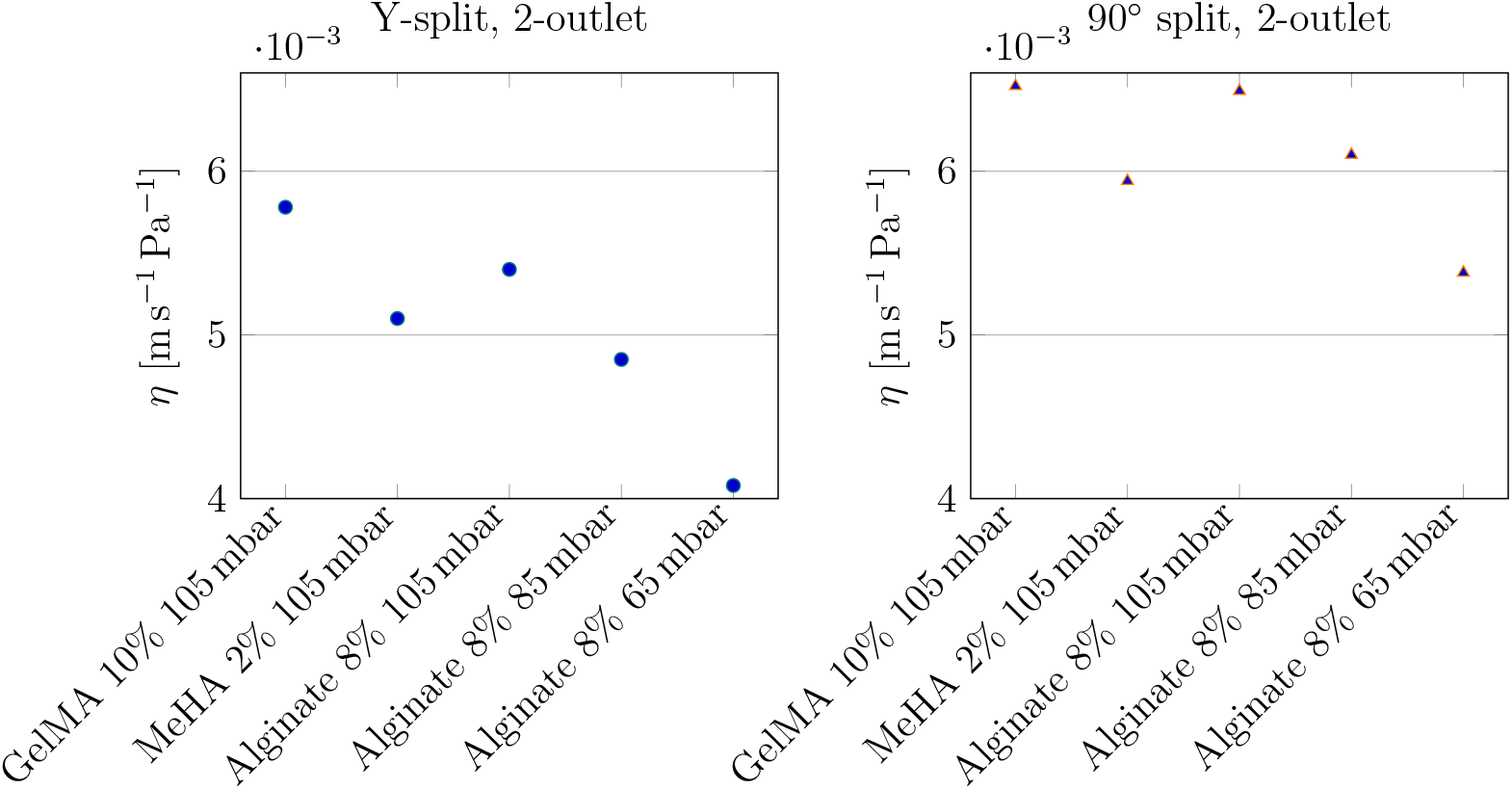
Efficiency index (pressure-normalised throughput) defined as mean outlet velocity divided by pressure drop, 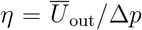, for 2-outlet Y-split and 90^°^ split designs across bioinks and inlet pressures.

Figure 9 presents a radar chart summarising five performance metrics, velocity, wall shear stress (WSS), efficiency, and outlet imbalance — for Alginate 8% at 105 mbar. The 90^°^ two-outlet configuration shows a balanced profile across all axes, indicating consistent performance with low shear and minimal imbalance. In contrast, the four-outlet Y-split achieves higher throughput (higher outlet velocity) but exhibits reduced pressure-normalised throughput and elevated shear, which may adversely affect cell viability.

**Figure 9:**
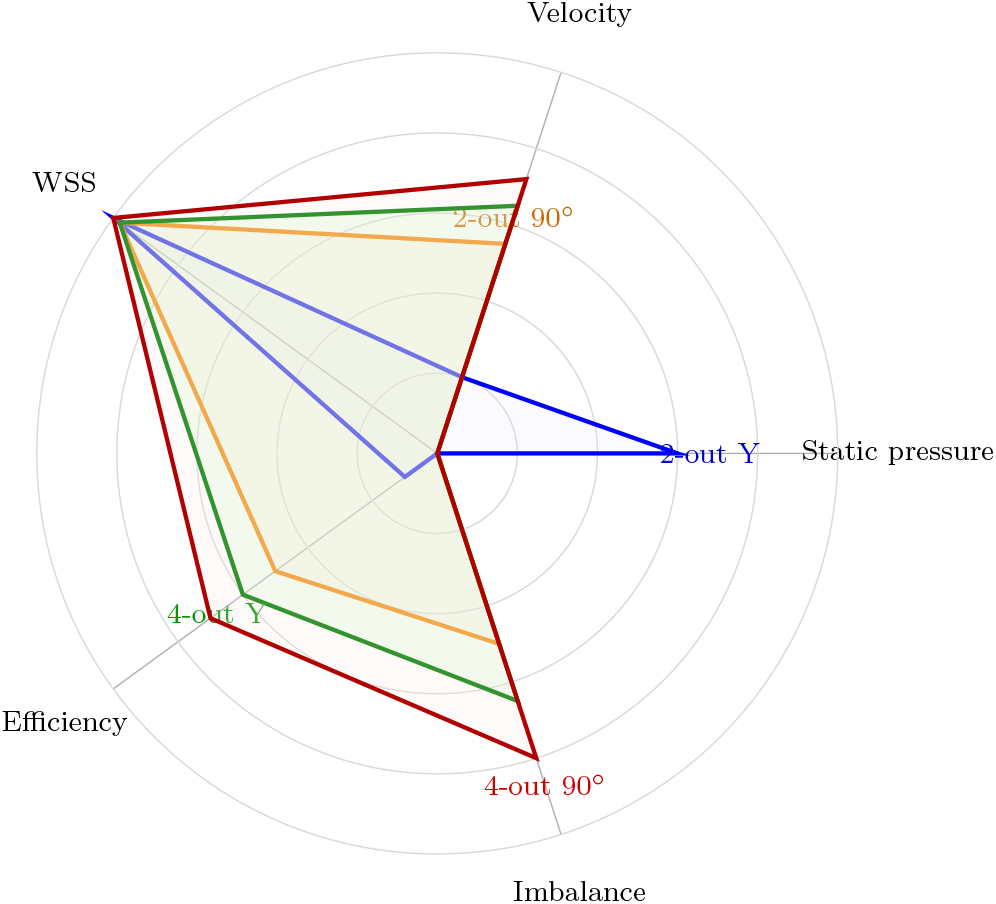
Radar plot comparing normalized values across five performance dimensions for 2- and 4-outlet Y-split and 90^°^ configurations using Alginate 8% at 105 mbar.

These multidimensional comparisons reinforce the need for task-specific nozzle selection rather than a single globally optimal design. For high-precision or viability-sensitive applications, minimising shear stress and outlet imbalance is the primary design priority. For rapid deposition or acellular support materials, higher velocity and pressure may be prioritised, justifying the use of geometries such as Y-splits despite their shear cost.

## 4 Discussion

Extrusion bioprinting performance is difficult to measure in narrow, opaque nozzles, optical velocimetry (PIV/µPIV) faces access and uncertainty limitations, particularly for scattering, cell-laden bioinks. As a result, CFD is routinely used to resolve pressure/WSS/velocity and compare candidate designs under controlled conditions (37; 22). Our simulations can already show that nozzle geometry strongly influences flow dynamics, stress distribution, and extrusion efficiency across a range of biomaterials and inlet pressures. We show that two-outlet nozzles exhibit superior flow symmetry, lower WSS, and higher efficiency than four-outlet counterparts, particularly under physiologically relevant printing conditions. Geometry and outlet size jointly tune WSS, pressure drop, and flow rate, which are the key determinants of viability and fidelity in extrusion bioprinting (22; 32). The 90^°^ 2-outlet design emerges as the most balanced configuration. As shown in Figures 4 and 6, it maintains low outlet flow imbalance (typically *I*_*Q*_ *<* 1%), moderate extrusion velocities, and the lowest WSS across all biomaterials tested. These attributes make it particularly suitable for applications involving cell-laden bioinks, where minimising mechanical stress is critical for preserving cell viability (46; 47; 10; 48). Mechanistically, cell survival decreases with increasing shear magnitude and cumulative exposure time. Nozzle size, pressure, extrusion speed, and bioink viscosity together set this mechanical burden (39). The preference for the 90^°^ 2-outlet design therefore aligns with reports that designs limiting cumulative high-shear exposure at matched flow targets better preserve viability (22). The efficiency index (Figure 8) further supports this design’s advantage by demonstrating superior pressure-normalised throughput at comparable operating pressures. In contrast, the Y-split 2-outlet configuration offers significantly higher outlet velocities, which may benefit applications requiring rapid deposition or high-throughput (e.g., printing of support matrices or acellular constructs). However, this advantage is offset by elevated WSS levels and a reduction in flow balance under high-viscosity and high-pressure conditions (Figure 7). The 31% imbalance observed in GelMA extrusion, for instance, can compromise print fidelity and uniformity in layer deposition, consistent with printability frameworks showing that excessive flow or poor rheology–process pairing degrades filament stability, filament–filament fusion, and pore geometry (32).

Transitioning to four-outlet designs introduces notable performance degradation. Both Y-split and 90^°^ geometries show reduced velocities and increased outlet imbalances (often exceeding 50%), particularly under low-pressure alginate conditions. These trends indicate that increasing outlet number imposes geometric and hydraulic penalties, likely stemming from accumulated minor losses and disrupted symmetry. While the 4-outlet 90^°^ design slightly outperforms its Y-split counterpart in flow uniformity and WSS containment, neither configuration matches the reliability of two-outlet designs, mirroring CFD observations that geometry-dependent penalties in WSS/flow must be weighed against the required throughput and biological tolerance (22; 49). The radar plot in Figure 9 consolidates these observations, highlighting that no single geometry dominates across all metrics. Instead, the trade-offs must be evaluated relative to application requirements.

Recent reviews explicitly encourage application-specific pairing of rheology, nozzle geometry, pressure, and speed to meet fidelity and viability targets—precisely the rationale motivating task-specific nozzles (32). Our findings thus advocate for a modular, task-specific nozzle strategy in bioprinting workflows, wherein geometry is selected dynamically based on bioink rheology, pressure constraints, and biological tolerance threshold (50; 51). They also suggest the value of active flow control mechanisms or real-time monitoring to mitigate imbalance and stress in high-output configurations, which may be further explored in future work. For cell-laden constructs, designs that minimize WSS and maintain flow uniformity should be prioritised. For acellular deposition, throughput becomes the primary consideration. *Overall, our results reinforce that nozzle design is not a universal choice but a task-dependent control knob that should be tuned explicitly to the dominant biological and manufacturing constraints of the intended print*. This aligns with reports that converging/tapered flow paths can reduce cumulative shear exposure for a given flow target, whereas more aggressive splits can increase stress and balance-control demands (22).

All simulations assumed steady, laminar flow under constant inlet pressure. This is appropriate for the low-Reynolds-number regime of hydrogel extrusion, but it does not capture transient effects during printing (pressure ramp-up, regulator fluctuations, stop–start toolpaths) that can briefly increase shear exposure and outlet imbalance. This work is also purely computational and lacks experimental validation. Fabricating the two- and four-outlet designs and performing simple extrusion tests (total flow rate and per-outlet collection at fixed pressures) would directly test the predicted balance and pressure–flow trends. Geometric dimensions were fixed to match one representative printer, and only 2 and 4 were simulated. Extending the same templates and pipeline to higher outlet counts would clarify when passive symmetry is insufficient. Finally, biological outcomes were inferred indirectly using wall shear stress as a proxy. Future work include incorporating residence-time estimates and basic post-extrusion viability assays to provide a direct mechanical-to-biological link. Overall, the qualitative ranking observed here is expected to remain unchanged, given the clear separations between designs and the symmetry-driven nature of the flow splitting. However, experimental validation is required to confirm absolute values and to assess ranking stability under practical operating conditions.

## 5 Conclusion

Task-specific multi-outlet nozzle designs were quantified using CFD simulations to quantify how multi–outlet nozzle geometry affects flow uniformity, WSS, and extrusion efficiency across representative bioinks and inlet pressures. The results show that nozzle geometry is a dominant factor governing extrusion performance. The two-outlet 90^°^ nozzle provides the best balance of flow symmetry, low WSS, and efficiency, indicating suitability for cell-laden hydrogel printing. The two-outlet Y-split achieves higher outlet velocities and throughput but incurs elevated shear, making it more appropriate for acellular or support materials. Increasing to four outlets consistently degraded balance and efficiency, highlighting the limits of passive scaling. Overall, these findings support our task-specific framework for selecting nozzle geometry based on biological and process constraints.

## CRediT authorship contribution statement

Conceptualization: MTB; Methodology: CDVU; Software/CFD: CDVU; Validation: CDVU; Formal analysis: CDVU; Investigation: CDVU and MTB; Resources: MTB; Data curation: CDVU; Writing—original draft: CDVU and MTB; Writing—review & editing: MTB; Visualization: CDVU; Supervision: MTB; Funding acquisition: MTB.

## Data and code availability

CSV metrics, CAD STEP/STL files, and plotting scripts will be provided as Supplementary Material.

## Acknowledgements

We acknowledge the University of Essex and the UC2 Lab for supporting this work.

## References

[1] S. V. Murphy and A. Atala, “3d bioprinting of tissues and organs,” Nature biotechnology, vol. 32, no. 8, pp. 773–785, 2014.

[2] C. Mandrycky, Z. Wang, K. Kim, and D.-H. Kim, “3d bioprinting for engineering complex tissues,” Biotechnology advances, vol. 34, no. 4, pp. 422–434, 2016.

[3] B. Yilmaz, A. Al Rashid, Y. Ait Mou, Z. Evis, and M. Koç, “Bioprinting: A review of processes, materials and applications,” Bioprinting, vol. 23, p. e00148, 2021.

[4] K. Hölzl, S. Lin, L. Tytgat, S. Van Vlierberghe, L. Gu, and A. Ovsianikov, “Bioink properties before, during and after 3d bioprinting,” Biofabrication, vol. 8, no. 3, p. 032002, 2016.

[5] B. Derby, “Printing and prototyping of tissues and scaffolds,” science, vol. 338, no. 6109, pp. 921–926, 2012.

[6] F. P. Melchels, J. Feijen, and D. W. Grijpma, “A review on stereolithography and its applications in biomedical engineering,” Biomaterials, vol. 31, no. 24, pp. 6121–6130, 2010.

[7] T. J. Hinton, Q. Jallerat, R. N. Palchesko, J. H. Park, M. S. Grodzicki, H.-J. Shue, M. H. Ramadan, A. R. Hudson, and A. W. Feinberg, “Three-dimensional printing of complex biological structures by freeform reversible embedding of suspended hydrogels,” Science advances, vol. 1, no. 9, p. e1500758, 2015.

[8] W. Sun, B. Starly, A. C. Daly, J. A. Burdick, J. Groll, G. Skeldon, W. Shu, Y. Sakai, M. Shinohara, M. Nishikawa et al., “The bioprinting roadmap,” Biofabrication, vol. 12, no. 2, p. 022002, 2020.

[9] X. Ma, M. Xu, X. Cui, J. Yin, and Q. Wu, “Hybrid 3d bioprinting of sustainable biomaterials for advanced multiscale tissue engineering,” Small, p. 2408947, 2025.

[10] B. Li, Z. Wang, C. Huang, L. Xu, S. Huang, M. Qu, Z. Xu, D. Zhang, B. Guo, T. Jin et al., “A comprehensive review on the printing efficiency, precision, and cell viability in 3d bioprinting,” Medical Engineering & Physics, p. 104448, 2025.

[11] E. Reina-Romo, S. Mandal, P. Amorim, V. Bloemen, E. Ferraris, and L. Geris, “Towards the experimentally-informed in silico nozzle design optimization for extrusion-based bioprinting of shear-thinning hydrogels,” Frontiers in bioengineering and biotechnology, vol. 9, p. 701778, 2021.

[12] J.C. Gómez-Blanco, J. B. Pagador, V.P. Galván-Chacón, L.F. Sánchez-Peralta, M. Matamoros, A. Marcos, and F.M. Sánchez-Margallo, “Computational simulation-based comparative analysis of standard 3d printing and conical nozzles for pneumatic and piston-driven bioprinting,” International Journal of Bioprinting, vol. 9, no. 4, p. 730, 2023.

[13] R. Gharraei, D. Bergstrom, and X. D. Chen, “Extrusion bioprinting from a fluid mechanics perspective,” 2024.

[14] C. Zhou, C. Liu, Z. Liao, Y. Pang, and W. Sun, “Ai for biofabrication,” Biofabrication, vol. 17, no. 1, p. 012004, 2024.

[15] J. C. G. Blanco, A. Macías-García, J.M. Rodríguez-Rego, L. Mendoza-Cerezo, F.M. Sánchez-Margallo, A. C. Marcos-Romero, and J. B. Pagador-Carrasco, “Optimising bioprinting nozzles through computational modelling and design of experiments,” Biomimetics, vol. 9, no. 8, p. 460, 2024.

[16] A. Malekpour and X. Chen, “Printability and cell viability in extrusion-based bioprinting from experimental, computational, and machine learning views,” Journal of functional biomaterials, vol. 13, no. 2, p. 40, 2022.

[17] U. N. M. Fareez, S. A. A. Naqvi, M. Mahmud, and M. Temirel, “Computational fluid dynamics (cfd) analysis of bioprinting,” Advanced Healthcare Materials, vol. 13, no. 20, p. 2400643, 2024.

[18] G. Ates and P. Bartolo, “Computational fluid dynamics for the optimization of internal bioprinting parameters and mixing conditions,” International Journal of Bioprinting, 2023.

[19] S. J. Müller, B. Fabry, and S. Gekle, “Predicting cell stress and strain during extrusion bioprinting,” Physical Review Applied, vol. 19, no. 6, p. 064061, 2023.

[20] Z. Guo, F. Fei, X. Song, and C. Zhou, “Analytical study and experimental verification of shear-thinning ink flow in direct ink writing process,” Journal of manufacturing science and engineering, vol. 145, no. 7, p. 071001, 2023.

[21] R. Sharma, I. P. Smits, L. De La Vega, C. Lee, and S. M. Willerth, “3d bioprinting pluripotent stem cell derived neural tissues using a novel fibrin bioink containing drug releasing microspheres,” Frontiers in bioengineering and biotechnology, vol. 8, p. 57, 2020.

[22] R. Chand, B. S. Muhire, and S. Vijayavenkataraman, “Computational fluid dynamics assessment of the effect of bioprinting parameters in extrusion bioprinting,” International Journal of Bioprinting, vol. 8, no. 2, p. 545, 2022.

[23] A. Blaeser, D. F. Duarte Campos, U. Puster, W. Richtering, M. M. Stevens, and H. Fischer, “Controlling shear stress in 3d bioprinting is a key factor to balance printing resolution and stem cell integrity,” Advanced healthcare materials, vol. 5, no. 3, pp. 326–333, 2016.

[24] J. Jia, D. J. Richards, S. Pollard, Y. Tan, J. Rodriguez, R. P. Visconti, T. C. Trusk, M. J. Yost, H. Yao, R. R. Markwald et al., “Engineering alginate as bioink for bioprinting,” Acta biomaterialia, vol. 10, no. 10, pp. 4323–4331, 2014.

[25] Y. B. Pottathara, M. Jordan, T. Gomboc, B. Kamenik, B. Vihar, V. Kokol, and M. Zadravec, “Solidification of gelatine hydrogels by using a cryoplatform and its validation through cfd approaches,” Gels, vol. 8, no. 6, p. 368, 2022.

[26] D. Petta, U. D’Amora, L. Ambrosio, D. W. Grijpma, D. Eglin, and M. D’Este, “Hyaluronic acid as a bioink for extrusion-based 3d printing,” Biofabrication, vol. 12, no. 3, p. 032001, 2020.

[27] T. Gregory, P. Benhal, A. Scutte, D. Quashie Jr., K. Harrison, C. Cargill, S. Grandison, M. J. Savitsky, S. Ramakrishnan, and J. Ali, “Rheological characterization of cell-laden alginate–gelatin hydrogels for 3d biofabrication,” Journal of the Mechanical Behavior of Biomedical Materials, vol. 136, p. 105474, 2022.

[28] L. Mendoza-Cerezo, J.M. Rodríguez-Rego, A. Macías-García, A. Callejas-Marín, L. Sánchez-Guardado, and A. C. Marcos-Romero, “Three-dimensional bioprinting of GelMA hydrogels with culture medium: Balancing printability, rheology and cell viability for tissue regeneration,” Polymers, vol. 16, no. 10, p. 1437, 2024.

[29] K. Yue, G. Trujillo-de Santiago, M. M. Alvarez, A. Tamayol, N. Annabi, and A. Khademhosseini, “Synthesis, properties, and biomedical applications of gelatin methacryloyl (gelma) hydrogels,” Biomaterials, vol. 73, pp. 254–271, 2015.

[30] N. Paxton, W. Smolan, T. Böck, F. Melchels, J. Groll, and T. Jungst, “Proposal to assess printability of bioinks for extrusion-based bioprinting and evaluation of rheological properties governing bioprintability,” Biofabrication, vol. 9, no. 4, p. 044107, 2017.

[31] C. B. Highley, C. B. Rodell, and J. A. Burdick, “Direct 3d printing of shear-thinning hydrogels into self-healing hydrogels.” Advanced Materials (Deerfield Beach, Fla.), vol. 27, no. 34, pp. 5075–5079, 2015.

[32] A. Schwab, R. Levato, M. D’este, S. Piluso, D. Eglin, and J. Malda, “Printability and shape fidelity of bioinks in 3d bioprinting,” Chemical reviews, vol. 120, no. 19, pp. 11 028–11 055, 2020.

[33] A. T. Banigo, L. Nauta, B. Zoetebier, and M. Karperien, “Hydrogel-based bioinks for coaxial and triaxial bioprinting: A review of material properties, printing techniques, and applications,” Polymers, vol. 17, no. 7, p. 917, 2025.

[34] P. J. McCauley, C. A. Fromen, and A. V. Bayles, “Cell viability in extrusion bioprinting: the impact of process parameters, bioink rheology, and cell mechanics,” Rheologica Acta, pp. 1–19, 2025.

[35] L. Lemarié, A. Anandan, E. Petiot, C. Marquette, and E.-J. Courtial, “Rheology, simulation and data analysis toward bioprinting cell viability awareness,” Bioprinting, vol. 21, p. e00119, 2021.

[36] S. Ramesh, O. L. Harrysson, P. K. Rao, A. Tamayol, D. R. Cormier, Y. Zhang, and I. V. Rivero, “Extrusion bioprinting: Recent progress, challenges, and future opportunities,” Bioprinting, vol. 21, p. e00116, 2021.

[37] M. R. Abdulwahab, Y. H. Ali, F. J. Habeeb, A. A. Borhana, A. M. Abdelrhman, and S. M. A. Al-Obaidi, “A review in particle image velocimetry techniques (developments and applications),” Journal of Advanced Research in Fluid Mechanics and Thermal Sciences, vol. 65, no. 2, pp. 213–229, 2020.

[38] A. Jahanbakhsh, K. L. Wlodarczyk, D. P. Hand, R. R. Maier, and M. M. Maroto-Valer, “Review of microfluidic devices and imaging techniques for fluid flow study in porous geomaterials,” Sensors, vol. 20, no. 14, p. 4030, 2020.

[39] H.-Q. Xu, J.-C. Liu, Z.-Y. Zhang, and C.-X. Xu, “A review on cell damage, viability, and functionality during 3d bioprinting,” Military Medical Research, vol. 9, no. 1, p. 70, 2022.

[40] Z. Lamberger, D. W. Schubert, M. Buechner, N. C. Cabezas, S. Schrüfer, N. Murenu, N. Schaefer, and G. Lang, “Advanced optical assessment and modeling of extrusion bioprinting,” Scientific Reports, vol. 14, no. 1, p. 13972, 2024.

[41] Ansys, Inc., Ansys Fluent User’s Guide, Release 2025 R1, Ansys, Inc., Canonsburg, PA, USA, 2025, fluent v25.1.

[42] G. Vivarelli, N. Qin, and S. Shahpar, “A review of mesh adaptation technology applied to computational fluid dynamics,” Fluids, vol. 10, no. 5, p. 129, 2025.

[43] M. S. Shakur, E. Lazarus, C. Wang, K. Du, I. V. Rivero, and S. Ramesh, “Effect of hydrodynamic shear stress on algal cell fate in 3d extrusion bioprinting,” Advanced Engineering Materials, vol. 27, no. 4, p. 2401768, 2025.

[44] J. Elango and C. Zamora-Ledezma, “Rheological, structural, and biological tradeoffs in bioink design for 3d bioprinting,” Gels, vol. 11, no. 8, p. 659, 2025.

[45] T. Suwannaphan, A. Kamnerdsook, S. Chalermwisutkul, B. Techaumnat, N. Damrongplasit, B. Traipattanakul, S. Kasetsirikul, and A. Pimpin, “Effects of shear and extensional stresses on cells: Investigation in a spiral microchannel and contraction– expansion arrays,” ACS Biomaterials Science & Engineering, 2025.

[46] Q. Wei, Y. An, X. Zhao, M. Li, J. Zhang, and N. Cui, “Optimal design of multi-biomaterials mixed extrusion nozzle for 3d bioprinting considering cell activity,” Virtual and Physical Prototyping, vol. 20, no. 1, p. e2438897, 2025. 10.36922/IJB025190182

[47] L. Lombardi, A. Scalzone, C. Ausilio, P. Gentile, and D. Tammaro, “Optimizing nozzle design in extrusion-based 3d bioprinting to minimize mechanical stress and enhance cell viability,” International Journal of Bioprinting, vol. 11, no. 4, pp. 315–327, 2025. [Online]. Available:

[48] S. Ahmad, H. Alam, and P. Thareja, “3d printing of hydrogels: a synergistic approach of rheology and computational fluid dynamics (cfd) modeling,” RSC Advances, vol. 15, no. 47, pp. 39 369–39 390, 2025.

[49] H. Ravanbakhsh, V. Karamzadeh, G. Bao, L. Mongeau, D. Juncker, and Y. S. Zhang, “Emerging technologies in multi-material bioprinting,” Advanced Materials, vol. 33, no. 49, p. 2104730, 2021.

[50] F. Chirianni, G. Vairo, and M. Marino, “Influence of extruder geometry and bio-ink type in extrusion-based bioprinting via an in silico design tool,” Meccanica, vol. 59, no. 8, pp. 1285–1299, 2024.

[51] M. Hidaka, M. Kojima, C. Zhang, Y. Okano, and S. Sakai, “Experimental and numerical approaches for optimizing conjunction area design to enhance switching efficiency in single-nozzle multi-ink bioprinting systems,” International Journal of Bioprinting, vol. 10, no. 5, p. 4091, 2024.

