## Supplementary Material for "Task-Specific Computational Fluid Dynamics Evaluation of Multi-Outlet Extrusion Nozzles for Bioprinting"

#### Supplementary File S1: Spatial field panels.

Table 5: Sensitivity of effective viscosity $\mu_{\text{eff}}$ evaluated at a representative shear rate of $\dot{\gamma} = 1000\ \text{s}^{-1}$. A $\pm 20\%$ variation in the consistency index $K$ and a $\pm 0.05$ variation in the flow behaviour index $n$ were tested for power-law bioinks. GelMA was modeled as Newtonian and therefore remains constant.

| **Bioink (model)** | **Baseline $\mu_{\text{eff}}$ [Pa·s]** | $K$ **−20%** | $K$ **+20%** | $n \pm 0.05$ |
|---|---|---|---|---|
| Alginate 8% (Power-law: $K = 56.9$, $n = 0.322$) | 0.526 | 0.421 (−20%) | 0.631 (+20%) | 0.372 (−29%) \| 0.743 (+41%) |
| MeHA 2% (Power-law: $K = 12.4$, $n = 0.55$) | 0.554 | 0.443 (−20%) | 0.665 (+20%) | 0.392 (−29%) \| 0.782 (+41%) |
| GelMA 10% (Newtonian: $\mu = 0.46$) | 0.460 | – | – | – |

Table 6: Geometry specification: 2-outlet 90° split nozzle (splitting block only; all dimensions in mm).

| Parameter | Value |
|---|---|
| Inlet diameter $D_{\text{in}}$ | 1.35 |
| Inlet straight length $L_{\text{in}}$ | 8.50 |
| Channel diameter $D_{\text{b}}$ | 1.35 |
| Straight run from junction to bend start $L_{\text{pre}}$ | 7.745 |
| Bend angle | 90° |
| Bend radius (centreline) $R_c$ | 1.00 |
| Straight run from bend end to outlet plane $L_{\text{post}}$ | 5.355 |
| Outlet diameter at split-block exit $D_{\text{out}}$ | 1.35 |

Table 7: Geometry specification: 2-outlet Y-split nozzle (splitting block only; all dimensions in mm).

| Parameter | Value |
|---|---|
| Inlet diameter $D_{\text{in}}$ | 1.35 |
| Inlet straight length $L_{\text{in}}$ | 8.50 |
| Channel diameter $D_{\text{b}}$ | 1.35 |
| Y-split bifurcation angle $\theta_Y$ | 30° |
| Branch angle relative to centreline | 15° |
| Branch centreline length (split to outlet plane) $L_{\text{branch}}$ | 15.611 |
| Branch diameter $D_{\text{branch}}$ | 1.35 |
| Outlet diameter at split-block exit $D_{\text{out}}$ | 1.35 |

Table 8: Geometry specification: 4-outlet 90° split nozzle (splitting block only; all dimensions in mm).

| Parameter | Value |
|---|---|
| Inlet diameter $D_{\text{in}}$ | 1.35 |
| Inlet straight length $L_{\text{in}}$ | 8.50 |
| Parent channel diameter $D_{\text{b1}}$ | 1.35 |
| *First split (parent → 2 branches)* | |
| Straight run from first split centre to bend start $L_{\text{pre},1}$ | 11.343 |
| Bend angle (tier 1) | 90° |
| Bend radius (centreline) $R_{c,1}$ | 1.00 |
| Bend centreline arc length (tier 1) $L_{\text{arc},1} = \frac{\pi}{2} R_{c,1}$ | 1.571 |
| Inter-tier straight length $L_{\text{inter}}$ | 3.00 |
| *Second split (each branch → 2 outlets)* | |
| Straight run from second split centre to bend start $L_{\text{pre},2}$ | 4.35 |
| Bend angle (tier 2) | 90° |
| Bend radius (centreline) $R_{c,2}$ | 1.00 |
| Bend centreline arc length (tier 2) $L_{\text{arc},2} = \frac{\pi}{2} R_{c,2}$ | 1.571 |
| Outlet diameter at split-block exit plane $D_{\text{out,SB}}$ | 1.35 |
| Outlet straight length inside split block $L_{\text{out,SB}}$ | 0.000 |
| Outlet plane used for reporting | Needle outlet plane (Table 10) |

Table 9: Geometry specification: 4-outlet Y-split nozzle (splitting block only; all dimensions in mm).

| Parameter | Value |
|---|---|
| Inlet diameter $D_{\text{in}}$ | 1.35 |
| Inlet straight length $L_{\text{in}}$ | 8.50 |
| Parent channel diameter $D_{\text{b1}}$ | 1.35 |
| *First split (parent → 2 branches)* | |
| Y-split bifurcation angle $\theta_{Y,1}$ | 60° |
| Branch angle relative to parent centreline | 30° |
| Centreline length (first split centre to next straight) $L_1$ | 15.015 |
| Branch diameter (tier 1) $D_{\text{b2}}$ | 1.35 |
| Inter-tier straight length $L_{\text{inter}}$ | 7.071 |
| *Second split (each branch → 2 outlets)* | |
| Y-split bifurcation angle $\theta_{Y,2}$ | 30° |
| Sub-branch angle relative to parent | 15° |
| Centreline length (second split centre to split-block exit plane) $L_2$ | 13.928 |
| Outlet diameter at split-block exit plane $D_{\text{out,SB}}$ | 1.35 |
| Outlet straight length inside split block $L_{\text{out,SB}}$ | 0.000 |
| Outlet plane used for reporting | Needle outlet plane (Table 10) |
| Outlet taper angle $\alpha$ | 0 (no taper) |
| Taper length $L_\alpha$ | – |

Table 10: Shared needle geometry used for all outlet channels (all dimensions in mm).

| Parameter | Value |
|---|---|
| Needle internal diameter at inlet $D_{\mathrm{needle,in}}$ | 1.35 |
| Needle internal diameter at outlet $D_{\mathrm{needle,out}}$ | 1.35 |
| Needle centreline length $L_{\mathrm{needle}}$ | 14.60 |
| Internal taper | None (constant internal diameter) |
| Taper length $L_{\alpha,\mathrm{needle}}$ | – |
| Taper half-angle $\alpha_{\mathrm{needle}}$ | – |
| Outlet plane used for reporting | Needle exit plane |

### Supplementary S2: Rheology fitting and viscosity bounds verification

#### Rheology fitting

To define extrusion-relevant rheology for the CFD simulations, literature-reported viscosity data were used to parameterise the constitutive models adopted for each bioink. For shear-thinning bioinks (8% alginate and 2% MeHA), apparent viscosity data were digitised from published flow curves and fitted using a power-law model. GelMA was modelled as a Newtonian fluid using a constant viscosity reported at printing temperature.

Figures S2–S4 show apparent viscosity as a function of strain rate on logarithmic axes, together with the fitted constitutive models. For alginate and MeHA, fitting was performed over the range $10^1$–$10^4$ s$^{-1}$, which spans the strain-rate regime relevant to extrusion through the studied nozzle geometries. The fitted parameters were used directly in all CFD simulations without further adjustment.

#### Shear-rate relevance

The strain-rate range used for rheological fitting was selected to reflect operating conditions within the nozzle geometries. Based on characteristic velocity and diameter scales, local strain rates within the split-block were estimated to lie primarily within the range $10^2$–$10^4$ s$^{-1}$, depending on inlet pressure and nozzle layout. The adopted fitting window therefore captures the shear regimes most relevant to extrusion through the studied geometries.

#### Viscosity bounds verification

At extremely low strain rates, shear-thinning power-law models predict unbounded viscosity as $\dot{\gamma} \to 0$. To maintain numerical stability in such regions, the CFD solver enforces effective viscosity limits internally. To verify that these limits did not distort the reported results, the minimum and maximum strain-rate magnitudes encountered within the computational domain were extracted from converged solutions, and the corresponding effective viscosities were evaluated using the adopted constitutive laws.

Table 11: Verification of effective viscosity limits for all simulated bioinks. Reported values correspond to the minimum and maximum strain-rate magnitudes encountered across all nozzle geometries and inlet pressures, together with the resulting effective viscosity range realised in the simulations.

| Bioink | $\dot{\gamma}_{\min}$ [s$^{-1}$] | $\dot{\gamma}_{\max}$ [s$^{-1}$] | $\mu_{\mathrm{eff}}$ range [Pa·s] | Limit active? |
|---|---|---|---|---|
| 8% Alginate | $9.82 \times 10^{-3}$ | $2.2 \times 10^{5}$ | 0.013–10 | Upper |
| 2% MeHA | $9.98 \times 10^{-3}$ | $3.17 \times 10^{5}$ | 0.040–10 | Upper |
| 10% GelMA | – | – | 0.46 | No |

In both shear-thinning cases, extremely low strain rates occur locally in near-stagnant regions, causing the effective viscosity to reach the solver-imposed upper limit. The reported viscosity ranges therefore reflect the bounded values realised during the simulations rather than the unbounded analytical limit of the power-law model.
